# *Pseudomonas aeruginosa* phenazines dictate site-specific competitive interactions with *Klebsiella pneumoniae*

**DOI:** 10.1101/2025.08.26.672413

**Authors:** Katlyn Todd, Olivia Schneider, Joshua M. Lawrence, Josefina L. Aronoff, Bartosz Witek, Valerie Velázquez-Colón, Verónica Santana-Ufret, Moraima Noda, Ryan F. Relich, Lifan Zeng, Dominique H. Limoli, Christopher Whidbey, Jay Vornhagen

## Abstract

*Pseudomonas aeruginosa* and *Klebsiella pneumoniae* are Gram-negative opportunistic pathogens that frequently colonize the human body and are major causes of infection. These bacteria are often co-isolated in polymicrobial urinary tract and lung infections, the latter of which is associated with increased disease severity and worse clinical outcomes. Despite their overlapping niches and clinical relevance, little is known about how these two pathogens interact and how those interactions influence human health. Given the growing recognition that microbial interactions are key drivers of disease, we investigated how *P. aeruginosa* and *K. pneumoniae* influence one another. We discovered an antagonistic interaction in which *P. aeruginosa* restricts the growth of *K. pneumoniae*. This inhibition is driven by phenazine production in *P. aeruginosa*, specifically the secondary metabolites pyocyanin and pyorubin, which are both necessary and sufficient to suppress *K. pneumoniae* growth. Using a diverse set of clinical isolates, we found that this antagonism is strain dependent. Both the susceptibility of *K. pneumoniae* to phenazines and the ability of *P. aeruginosa* to restrict *K. pneumoniae* growth varies between strains. Moreover, the necessity of phenazine production is specific to the site of infection. Together, these findings demonstrate that strain background and environmental context are critical determinants of pathogen interactions. Our work underscores the importance of considering these variables when investigating how microbial interactions influence infection and disease outcomes.

## Introduction

Microbial interactions are increasingly recognized as critical determinants of human health, reflected by the rapid growth of microbiome research. These interactions influence a wide range of physiological outcomes and contribute to human development and disease (conceptually reviewed in [1]). The gut is a well-studied site where trillions of viruses, bacteria, and microbial eukaryotes shape immunological development, metabolism, disease susceptibility, endocrine function, and neurological health (reviewed in [2–6], amongst many others). Beyond the gut, microbial interactions influence host biology in the oral cavity, lung, nasopharynx, respiratory tract, skin, and urinary and reproductive systems. Yet, little is known about the specific mechanisms that underpin these interactions, and how they may shape human health.

*Pseudomonas aeruginosa* (Pa) and *Klebsiella pneumoniae* (Kp) are Gram-negative opportunistic pathogens that are frequent colonizers of the human body (reviewed in [7–9]). They are also important causes of similar bacterial infections, such as pneumonia, urinary tract infections (UTI), and bacteremia. Despite occupying similar niches, little is known about how these bacteria interact. Some laboratory-based studies suggest that Pa can restrict Kp growth [10]. *In vivo* studies and clinical reports indicate that concurrent infection is associated with worse outcomes [11–14]. Commensurately, other laboratory-based studies demonstrate that Pa and Kp can co-exist in the same niche [15–17]. This niche overlap implies that interaction between these genera is possible, which is supported by evidence of genetic transfer between them [18].

Understanding the molecular interactions between Pa and Kp is important for several reasons. First, both can be highly resistant to antibiotics, which poses significant threats to public health. Antibiotic resistance greatly complicates infection treatment, leading to extreme excess healthcare costs and mortality. Kp and Pa are ranked as top ten 2024 WHO Bacterial Priority Pathogens, and both organisms are ranked in the top six most important causes of healthcare burden due to antimicrobial resistance [19, 20]. Second, Pa and Kp employ distinct strategies to maximize fitness in polymicrobial environments. Pa is highly responsive to microbial and environmental cues, deploying quorum sensing (QS) networks (reviewed in [21]), secreted and contact-dependent antagonistic mechanisms (for example [22–24]), and forming complex biofilms that enhance environmental adaptability (reviewed in [25, 26]). In contrast, Kp utilizes metabolic flexibility and stress resistance to persist in diverse niches (reviewed in [26, 27]). Additionally, the molecular mechanisms underlying host-specific fitness likely differ between these genera (reviewed in [26]). Third, Pa and Kp are highly responsive to environmental context (for example [28–30]), and possess extensive genetic diversity, with both genera harboring open pangenomes exceeding 100,000 genes. Their pangenomes are comprised of conserved core genes and variable accessory elements which confer adaptability and strain-specific phenotypes. This genetic plasticity shapes behaviors like nutrient acquisition, resilience to environmental stress, and the deployment of antagonistic effectors (reviewed in [9, 31, 32]). Finally, metabolic environment greatly impacts Kp and Pa pathogenesis [33–36]. As such, studying how Pa and Kp interact is important for understanding microbial interactions in general, and how they apply to human health.

This study is built off an initial observation that mice colonized with native Pa strains demonstrated low Kp gut colonization density. Thus, we hypothesized that Pa directly restricts Kp growth. We aimed to determine 1) if Pa can restrict Kp growth, 2) if restriction was due to overlapping nutritional niches, contact-dependent killing, or production of Kp-restrictive secondary metabolites, and 3) what pathways are required for Pa restriction of Kp growth. The work presented here demonstrates that Pa-Kp competition is multifactorial but primarily driven by production of Kp-restrictive secondary metabolites, which is dependent both on strain and environmental context.

## Results

In the process of performing *in vivo* experiments unrelated to this study, we observed the presence of an indigenous gut bacterium resistant to kanamycin and rifampicin. This bacterium was associated with low Kp colonization 24 to 48 hours after oral inoculation (**Figure S1A**). We isolated these strains of this bacterium, which we named 145.1, 191.1, and 193.1 based on the mouse of origin, and identified them as Pa based on morphology and matrix-assisted laser desorption ionization–time of flight (MALDI-TOF) mass spectrometry. Whole-genome sequencing of these strains revealed that they were most similar to sequence type ST175, and using a subset of Pa isolates from a previous study [37], we determined that these strains grouped with Clade A (PAO1 major group, **Figure S1B**).

To determine if these wild Pa were able to restrict Kp growth, potentially explaining the observed reduction in Kp gut colonization density, we performed simple co-culture assays in Luria-Bertani (LB) broth. The growth of Kp strain KPPR1 was restricted 99.4%, 98.7%, and 97.5% by Pa 145.1, 191.1, and 193.1, respectively (**Figure 1A**). Given that the behavior of Pa is highly strain-dependent, we aimed to contextualize our finding using two well-characterized lab Pa strains, PAO1 and PA14. PAO1 exhibited comparatively less restriction despite its similarity of the wild Pa strains, only inhibiting 76.4% of KPPR1 growth (**Figure 1B**), whereas PA14 was highly restrictive, inhibiting 99.3% of KPPR1 growth (**Figure 1C**). Next, we sought to determine if this phenotype was contact-dependent. To this end, we grew KPPR1 in filter-sterilized spent media of each Pa strain, using KPPR1 self-spent media as a control. As expected, KPPR1 growth was highly restricted in its own sterile spent media, achieving an average of 1.2% of its growth in fresh LB broth (**Figure 1D-F**). Interestingly, KPPR1 growth was significantly more restricted in Pa 145.1-, 191.1-, and 193.1-spent LB broth (70.7%, 99.0%, and 98.5% restriction of growth compared to self-spent media, respectively, **Figure 1D**).

**Figure 1.**
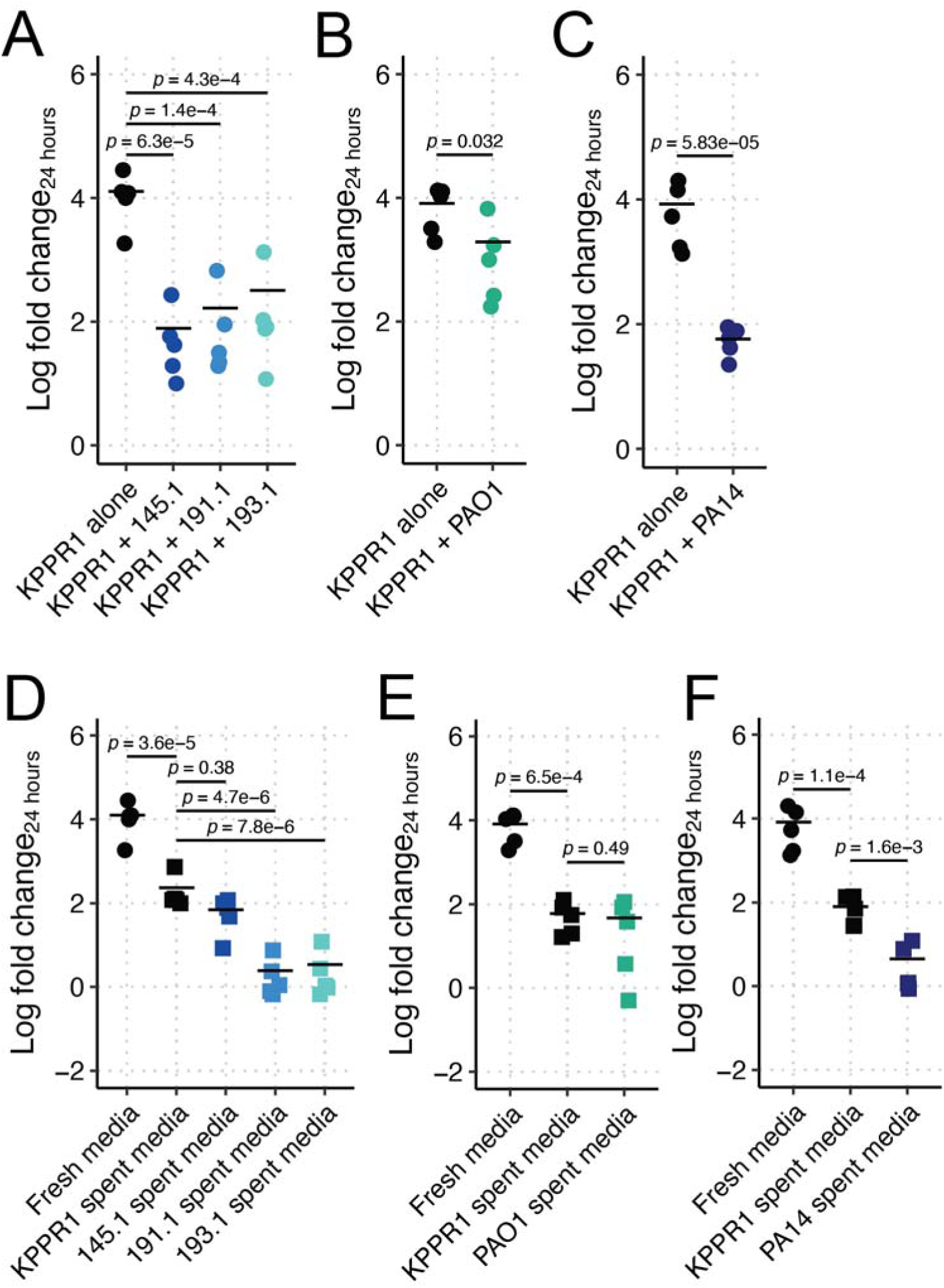
Pa restricts Kp growth in LB in a strain-dependent, contact-independent manner. KPPR1 was grown alone or in co-culture in LB with mouse-derived wild Pa (**A**), PAO1 (**B**), PA14 (**C**) or in filter sterilized spent media of KPPR1 or each Pa strain (**D-F**). For **A-F**, “Log fold change_24 hours_” = log_10_(output KPPR1 CFU at 24 hours/input KPPR1 CFU). *p-*values represent Tukey multiple comparison correction following one-way ANOVA. Each data point is a biological replicate, and horizontal lines indicate the mean of each dataset.

To contextualize these findings with PAO1 and PA14, we repeated these assays using these strains. Similar to co-culture results, PAO1-spent media did not significantly restrict KPPR1 growth (21.0% restriction of growth compared to self-spent media, **Figure 1E**), but PA14-spent media was highly restrictive (94.2% restriction of growth compared to self-spent media, **Figure 1F**). Collectively, these data indicate that Pa restriction of Kp growth is contact-independent, but strain-dependent.

Next, we aimed to assess the specificity of the Kp growth restriction phenotype to Pa. First, we performed co-culture competitions with the *Escherichia coli* (Ec) strain MG1655. Notably, Ec and Kp have similar nutritional niches, which has been exploited to reduce Kp gut colonization in model colonization systems [38, 39]. Ec MG1655 was unable to restrict KPPR1 (9.5% inhibition, **Figure S2A**). Then, we tested our approach using the Kp strain 13F11, which is a KPPR1 mutant with an intergenic mariner transposon insertion. We reasoned that this strain should not restrict KPPR1 growth as they are nearly identical strains. As expected, Kp 13F11 did not significantly restrict KPPR1 growth (25.6% inhibition, **Figure S2B**). Neither Ec MG1655 nor Kp 13F11 spent media significantly restricted growth of KPPR1 beyond self-spent media (52.6% and 12.1% restriction of growth compared to self-spent media, respectively, **Figure S2C-D**). Next, we assayed the dependency of this phenotype on growth medium. To this end, we repeated our competition assays in M9 minimal medium supplemented with 1.0% casamino acids (“M9-cas”). Pa restriction of KPPR1 growth in M9-cas was comparable to LB broth (**Figure S3**), and Ec MG1655 and Kp 13F11 were similarly unable to restrict KPPR1 growth (**Figure S4**). These data indicate that growth restriction of KPPR1 was limited to Pa and was insensitive to the media we tested.

Given that M9-cas and LB broth have similar available carbon sources, we hypothesized that the wild Pa strains and PA14 may be exhausting specific nutrients from these media more efficiently than that of Ec MG1655, Kp 13F11, or Pa PAO1. First, we measured the growth of these strains in both media types. KPPR1 had an early growth advantage over all Pa strains in LB broth that was mitigated by the end of the experiment, except for PA14, which reached a slightly lower final density than KPPR1 (**Figure S5A-B**). Ec MG1655 demonstrated a slight growth deficit compared to KPPR1, and as expected, Kp 13F11 growth was identical to KPPR1 (**Figure S5C-D**). Pa growth results in M9-cas were different than that of LB broth, wherein Pa grew to higher densities than KPPR1 (**Figure S5F-G**). Ec MG1655 and Kp 13F11 growth was identical to KPPR1 in M9-cas (**Figure S5H-I**). Given that Pa restriction of Kp was observed in both media, we concluded that the growth restriction phenotype was independent of growth characteristics, as Pa outgrew KPPR1 in M9-cas, but not LB broth.

Next, we aimed to determine if the metabolic potential of Pa and KPPR1 was similar. To this end, we assayed all five Pa strains, Ec MG1655, and KPPR1 for their ability to utilize different carbon sources using the BioLog PM1 and PM2 plates. No single carbon source was utilized more efficiently by the inhibitory Pa strains (PA14, 145.1, 191.1, 193.1) than the non-inhibitory strain PAO1 or KPPR1 (**Figure S6A**). Additionally, the Pa strains demonstrated a distinct carbon utilization profile compared to KPPR1 (**Figure S6B**), suggesting that differences in carbon source utilization were not explanatory for the restriction phenotype. Results with non-inhibitory Ec MG1655 were similar to the inhibitory Pa strains (**Figure S6**), further indicating that differences in carbon source utilization did not explain Kp growth restriction.

Despite having distinct carbon utilization profiles, we did not exclude the possibility that Pa is exhausting specific nutrients from both LB broth and M9-cas. Thus, we reasoned that nutrient supplementation could fully restore KPPR1 growth. We tested the ability of KPPR1 to grow in spent LB broth supplemented with casamino acids. We observed that growth was fully restored when self-spent LB broth was supplemented with casamino acids (213.7-fold increase in growth over water supplementation, **Figure 2A**). Interestingly, casamino acid supplementation was insufficient to restore KPPR1 growth to fresh LB broth levels in 145.1-, 191.1-, and 193.1-spent LB broth, or to the degree that supplementation restored growth in self-spent LB broth (21.6, 18.6, and 104.9-fold increase in growth over water supplementation, **Figure 2A**). Similar results were observed for PAO1 and PA14; however, casamino acid supplementation not only failed to restore growth in either PAO1 or PA14 spent LB broth but also failed to significantly enhance KPPR1 growth beyond PAO1 or PA14 spent LB broth (**Figure 2B**).

**Figure 2.**
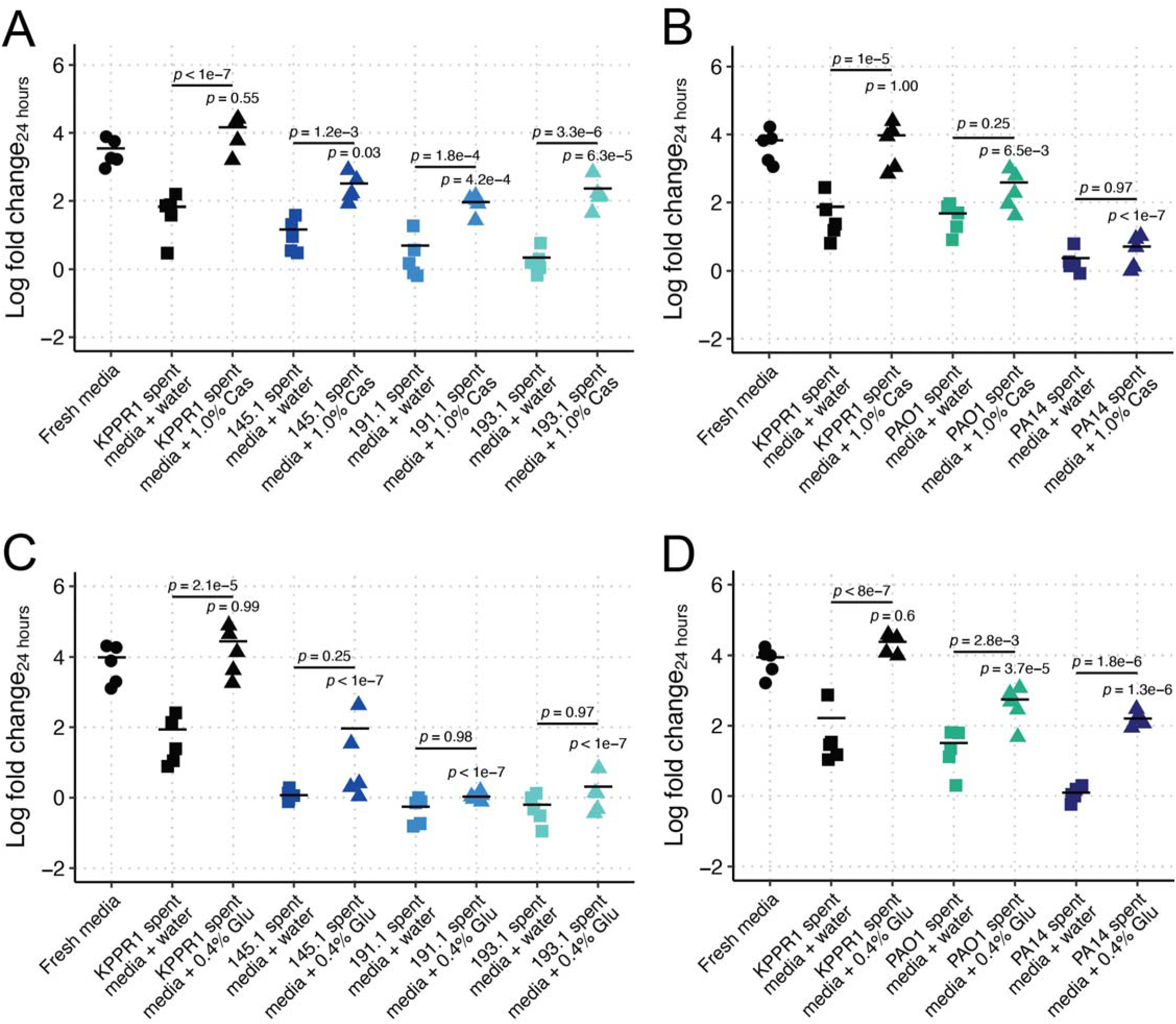
Casamino acid and glucose supplementation are insufficient to complement Kp growth restriction in Pa-spent LB. KPPR1 was grown alone or in filter sterilized spent LB media of KPPR1 or mouse-derived wild Pa (**A, C**) or PAO1 and PA14 (**B, D**) supplemented with water, 1% casamino acids (“Cas,” **A-B**) or 0.4% glucose (“Glu,” **C-D**). For **A-D**, “Log fold change_24 hours_” = log_10_(output KPPR1 CFU at 24 hours/input KPPR1 CFU). *p-*values represent Tukey multiple comparison correction following one-way ANOVA. *p-*values over columns indicate comparison to “Fresh media” condition. Each data point is a biological replicate, and horizontal lines indicate the mean of each dataset.

Finally, we tested if we were able to completely restore KPPR1 growth in Pa-spent LB broth by supplementing a standard carbon and energy source: glucose. Supplementation of glucose to self-spent LB broth fully restored KPPR1 growth (326.5-fold increase growth over water supplementation, **Figure 2C**). Like casamino acid supplementation, glucose supplementation of Pa 145.1-, 191.1-, and 193.1-spent LB broth was insufficient to restore Kp KPPR1 growth relative to fresh LB or glucose-supplemented self-spent LB broth (11.5, 1.9, and 3.2-fold increase over water supplementation, **Figure 2C**). Glucose supplementation was more restorative of Kp growth in PAO1 and PA14 than casamino acids; however, KPPR1 growth was still not restored to that of fresh media levels (**Figure 2D**). Collectively, these data suggest that an overlapping nutritional niche between Kp and Pa does not fully explain the restriction of Kp growth.

Given that Pa inhibition of KPPR1 growth is contact-independent and is at least partially independent of the nutritional niche, we reasoned that there is a factor or set of factors that Pa secretes to inhibit Kp growth. To identify candidate factors, we screened the PA14NR transposon library. To this end, we generated a GFP-expressing KPPR1 strain using the pJL1-sfGFP plasmid and screened 5,708 Pa transposon mutants for their ability to restrict this strain, using fluorescent intensity as a proxy for KPPR1 growth (**Figure S7A**, **Table S1**). This screen was repeated twice, and PA14 transposon mutants from co-cultures that had a fluorescent intensity |z-score| > 2.5 in both replicates were selected for validation. This yielded 18 candidate factors, 16 of which were unable to fully restrict KPPR1 growth and two that highly restricted KPPR1 growth.

The 18 transposon mutants identified as candidate factors involved in KPPR1 growth restriction were then validated for their ability to restrict KPPR1 growth. First, we determined if any of the non-restrictive phenotypes we observed were due to growth defects. Only one mutant, *tpiA* (PA14_62830), demonstrated a baseline growth defect, although this defect was modest (**Figure S7B-C**). Next, we tested the 18 mutants for their ability to restrict KPPR1 growth in co-culture and spent media assays (**Figure S7D-E**). 7/18 mutants displayed a phenotype analogous to our original screen in either the co-culture or spent media culture. Thus, this screen yielded 7 genes involved in Pa restriction of Kp growth (**Table 1**).

**Table 1.** Validated factors involved in KPPR1 growth restriction.

| PA14 gene locus | PAO1 gene locus | Gene name | Gene description | z-score |
| --- | --- | --- | --- | --- |
| PA14_01960 | PAO157 |  | putative RND efflux membrane fusion protein precursor | 4.03 |
| PA14_06570 | PA0504 | *bioD* | dethiobiotin synthase | 6.52 |
| PA14_18520 | PA3543 | *algK* | alginate biosynthetic protein AlgK precursor | 3.03 |
| PA14_19120 | PA3477 | *rhlR* | acylhomoserine lactone dependent transcriptional regulator | 4.14 |
| PA14_32750 | PA2465 |  | hypothetical protein | 3.3 |
| PA14_51430 | PA0996 | *pqsA* | probable coenzyme A ligase | 2.9 |
| PA14_62830 | PA4748 | *tpiA* | triosephosphate isomerase | 3.05 |

We observed that several mutants in our screen appeared to have modified phenazine production. The *psqA*, *rhlR*, and *tpiA* mutants did not produce high amounts of pyocyanin (PYO) and pyorubin (PYR). PYR collectively refers to 5-methylphenazine-1-carboxylate (5MPCA) and its spontaneously synthesized derivates including 5-methyl-7-amino-1-carboxyphenazinium betaine (Aeruginosin A) and 5-methyl-7-amino-1-carboxy-3-sulfophenazinium betaine (Aeruginosin B) [40, 41]. Production of blue PYO and red PYR lead to coloration of spent culture media when in high enough concentrations. Both PYO and PYR have previously been reported to have antimicrobial activities [42, 43]. Thus, we aimed to test their role in Kp restriction. First, we confirmed our screen results using marker-less mutants to ensure that we were not observing unexpected effects of transposon mutagenesis. RhlR binds the autoinducer *N*-butanoyl-L-homoserine lactone, which is produced by RhlI, leading to expression of many genes, including the phenazine biosynthesis loci [44, 45]. Both the marker-less Δ*rhlR* and Δ*rhlI* mutants were unable to restrict KPPR1 growth in our co-culture and spent media assays, (**Figure 3A-B**) in both LB broth (**Figure 3A-B**), and M9-cas (**Figure S8A-B**) compared to their parental strain, PA14. Consistent with WT PA14 results (**Figure 2B, D**), supplementation of Δ*rhlR* and Δ*rhlI* spent media with casamino acids was insufficient to restore KPPR1 growth (**Figure S8C**), whereas glucose supplementation fully restored KPPR1 growth (**Figure S8D**). Importantly, the Δ*rhlR* and Δ*rhlI* mutant strains grow equivalently to PA14 in both LB and M9-cas (**Figures S5E, S5J**); thus, this phenotype is dependent on products of the RhlRI regulon, which includes phenazines.

**Figure 3.**
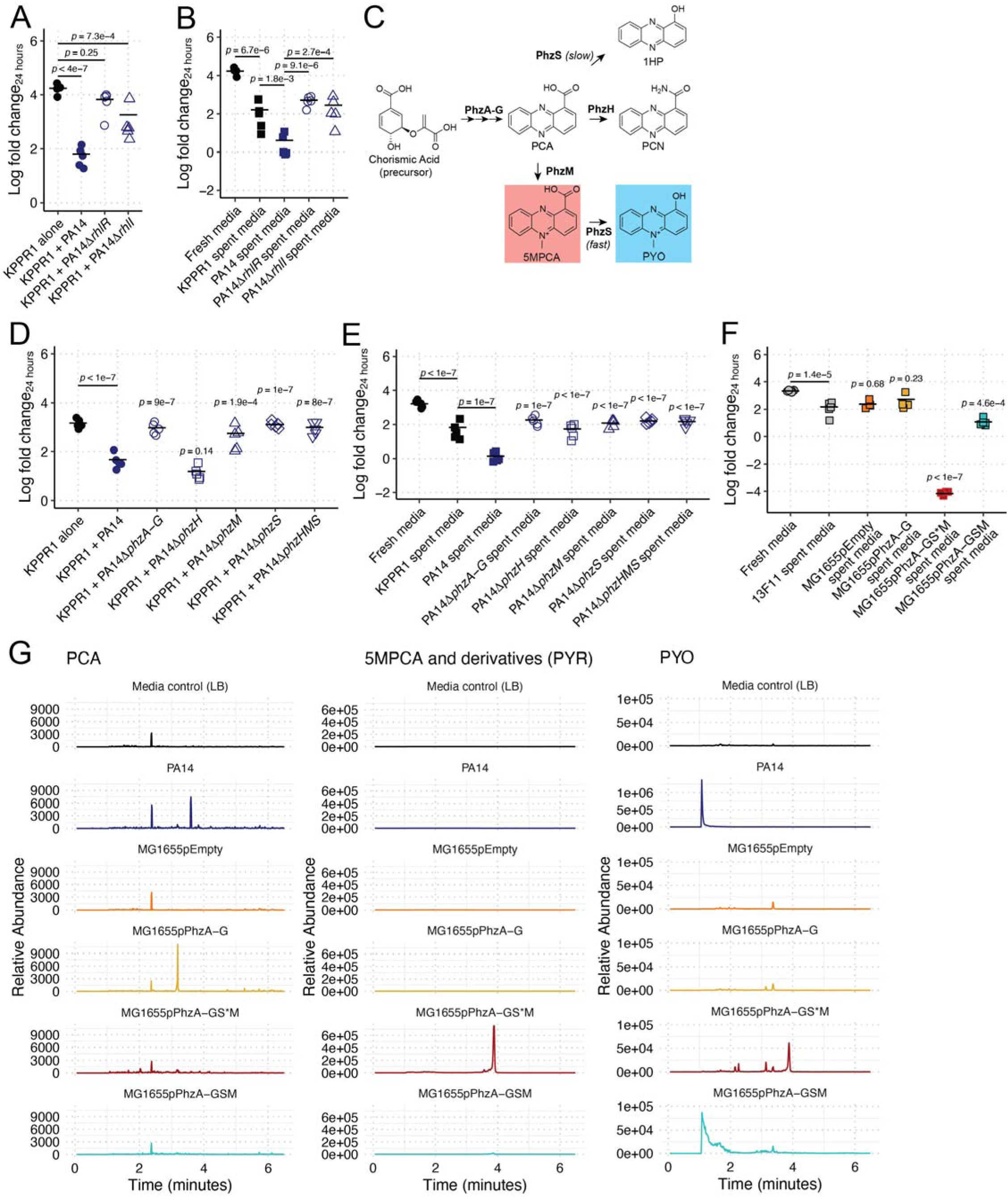
Phenazines are necessary and sufficient for Kp growth restriction. KPPR1 was grown alone or in co-culture in LB with WT PA14, PA14Δ*rhlR*, or PA14Δ*rhlI* (**A**) or in filter sterilized spent media of KPPR1 or each Pa strain (**B**). Phenazine biosynthesis (**C**) is under the transcriptional control of RhlIR. KPPR1 was grown alone or in co-culture in LB with WT PA14, PA14Δ*phzA-G*, PA14Δ*phzH*, PA14Δ*phzM*, PA14Δ*phzS*, or PA14Δ*phzHMS* (**D**) or in filter sterilized spent media of KPPR1 or each Pa strain (**E**). Additionally, 13F11 (Kan^R^ KPPR1 variant) was grown in filter sterilized spent media of 13F11 or MG1655 containing an empty vector (pEmpty) or constitutively expressing PCA (pPhzA-G), PYR (pPhzA-GS*M), and PYO (pPhzA-GSM, **F**). Datapoints outlined in red are below the limit of detection (200 CFU/mL). For **A-B** and **D-F**, “Log fold change_24 hours_” = log_10_(output KPPR1 CFU at 24 hours/input KPPR1 CFU). Each data point is a biological replicate, and horizontal lines indicate the mean of each dataset. *p-*values represent Tukey multiple comparison correction following one-way ANOVA. For **D**, **E**, and **F**, *p-*values above individual columns indicate comparison to “KPPR1 + PA14,” “PA14 spent media,” and “13F11 spent media” groups, respectively. LC-MS was used to quantify phenazine secretion from Ec and Pa strains (**G**). Note that the scale for the PA14 PYO chromograph is larger than the other PYO chromographs to accommodate the large peak. PA14 and MG1655 pPhzA-GSM chromographs on the same scale can be found in **Figure S14B**.

Then, we repeated our co-culture and spent media assays with strains containing deletions in the *phzA-G*, *phzH*, *phzM*, *phzS*, and *phzHMS* loci. PhzA-G are necessary to synthesis phenazine-1-carboxylic acid (PCA), which is converted to 5MPCA by PhzM and from 5MPCA to PYO by PhzS or directly converted from PCA to 1-hydroxyphenazine (1HP) by PhzS [46]. Additionally, PhzH converts PCA to phenazine-1-carboxamide (PCN). Thus, deletion of *phzA-G* renders Pa unable to synthesize PCA, and thus PYO, 5MPCA (and thus, PYR), 1HP, or PCN. Deletion of *phzH* ablates PCN production, deletion of *phzS* ablates PYO and 1HP production, deletion of *phzM* ablates 5MPCA and PYO, and deletion of *phzHMS* ablates production of PYO, PYR, 1HP, and PCN but maintains PCA production (**Figure 3C**).

Interestingly, we observed significantly less Kp growth restriction by the Δ*phzA-G*, Δ*phzM*, Δ*phzS*, and Δ*phzHMS* mutants in co-culture, and no impact with the Δ*phzH* mutant (**Figure 3D**). Similar results were observed in our spent media assay, though we did observe significantly less growth restriction in Δ*phzH*-spent LB broth (**Figure 3E**). Collectively, this suggests that the phenazine PYO and/or PYR are necessary for the observed growth restriction phenotype.

Most studies investigating the antimicrobial mechanisms of phenazines have focused on PYO. Due to the chemical similarity between PYO and other phenazines, it is commonly assumed that these compounds exert similar biological effects. The prevailing model for PYO’s antimicrobial activity involves the induction of oxidative stress: PYO readily diffuses across cellular membranes and promotes the formation of superoxide radicals, leading to cellular toxicity (reviewed in [47]). Thus, to determine if PYO is sufficient for restriction of Kp growth, we used a heterologous expression system in Ec MG1655 to constitutively produce PCA, PYR, and PYO, testing Kp (strain 13F11, Kan^R^ KPPR1 strain) in the resulting spent media. As expected, the empty vector and PCA over-producing strains failed to inhibit Kp growth beyond that of self-spent media (**Figure 3F**). Conversely, the spent media from the PYO over-producing strain inhibited Kp growth (**Figure 3F**) to a similar degree as WT PA14 (**Figure 1F**), demonstrating that PYO is sufficient to restrict Kp growth.

Surprisingly, the spent media from the PYR over-producing strain was completely bactericidal (**Figure 3F**). Of note, PYR overproduction was a result of a spontaneous frameshift mutation at T998 of *phzS* (annotated as “pPhzA-GS*M”) that ablated the final step of PYO production, leading to rapid accumulation 5MPCA and its derivates. High-resolution mass spectrometry of the spent medium of this strain confirmed the presence of 5MPCA and that it was absent in the spent media of other heterologous expression strains (**Figure 3G**). A previous report of a 5MPCA-sensitive microbe indicated that the activity of 5MPCA is dependent on its oxidation state, wherein its reduced form is more active than its oxidized form [42]. To determine if this is true for the activity of PYR against Kp, we titrated a reducing agent into spent media from our PYR producing Ec strain and measured its ability to kill Kp. Consistent with previous reports, the addition of reducing agent (generating reduced 5MPCA) is more active against Kp (**Figure S9**). Taken together, these data show that phenazines are both necessary and sufficient to restrict Kp growth.

Next, we aimed to determine the minimum inhibitory (MIC) and bactericidal (MBC) concentrations of PYO and PYR for KPPR1. To replicate our co-culture and spent media culture assays, we measured the MIC and MBC of PYO in both fresh LB and PA14Δ*phzA-G* spent LB. The MIC of PYO was 8-and 32-fold lower than the MBC (**Figure 4A-B**), in fresh and spent media, respectively, indicating that PYO is primarily bacteriostatic, which is consistent with the ability of PYO to arrest bacterial respiration [48], as opposed to kanamycin, which has an MIC only 2-fold lower than the MBC, indicating bactericidal activity (**Figure 4A-B**). We determined that PYR is primarily bactericidal, as the MIC is at most 2-fold lower (**Figure 4C**) than the concentration where bactericidal activity was detected (**Figure 3F**). Interestingly, PYO and PYR display concentration-dependent additive (0.5 < Fractional Inhibitory Concentration < 4, % MG1655pPhzA-GS*M spent media = 3.13%) and synergistic effects (Fractional Inhibitory Concentration > 0.5, % MG1655pPhzA-GS*M spent media = 6.25%, 12.5%, 25%, **Figure 4D**).

**Figure 4.**
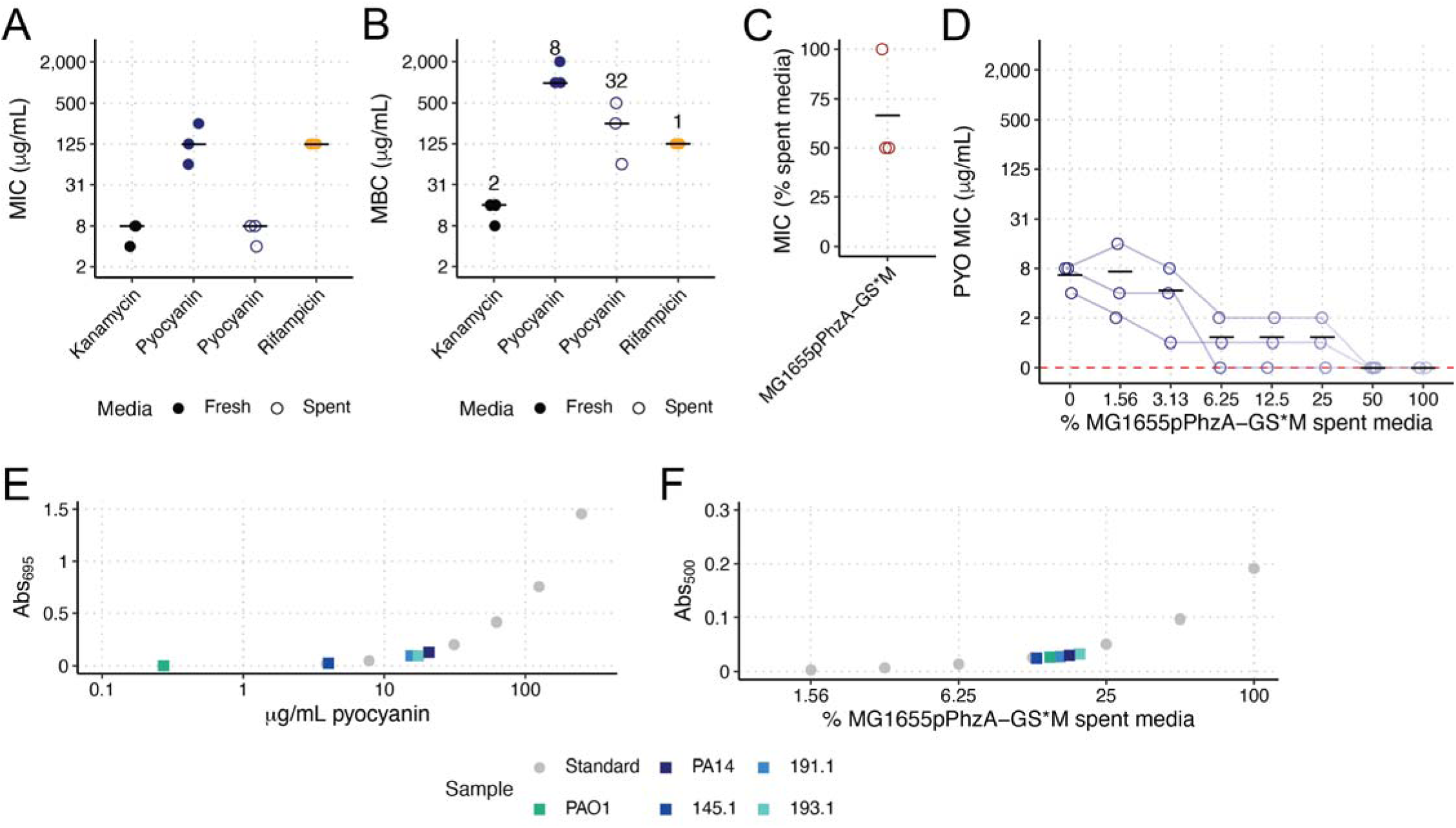
Minimum inhibitory and bactericidal concentrations of phenazines. The minimum inhibitory concentration (MIC) and minimum bactericidal concentrations (MBC) were determined for KPPR1 for kanamycin, to which KPPR1 is sensitive, PYO, and rifampicin, to which KPPR1 is resistant (**A-B**). The numbers above each column in **B** represent the median MBC:MIC ratio. The MIC of PYR (MG1655pPhzA-GS*M spent media) was also calculated (**C**). The effect of 5MPCA concentration on the PYO MIC was also measured (**D**). Each data point is a biological replicate, and horizontal lines indicate the mean of each dataset. The Abs_695_ and Abs_500_ of filter sterilized spent LB from mouse-derived wild Pa, PAO1, and PA14 were measured and PYO concentrations were interpolated from a PYO (**E**) and PYR (**F**) standard curve. For **E-F**, only median values of three biological replicates are shown.

Finally, we estimated the amount of PYO and PYR for each of our WT Pa strains produced in conditions identical to our spent media assay. PYO results were concordant with our co-culture and spent media assays, wherein PAO1 produced the least PYO (0.272 mg/mL) and was the least restrictive strain, followed by 145.1 (4.02 mg/mL), 191.1 (15.5 mg/mL), 193.1 (17.4 mg/mL), and PA14 (20.7 mg/mL, **Figure 4E**). All five strains displayed little PYR production (**Figure 4F**), measured at Abs_500_ (**Figure S10A**), concordant with high-resolution mass spectrometry for PA14, which displayed no 5MPCA production (**Figure 3G**). Only 193.1 displayed an Abs_500_ higher than PA14, suggesting limits to the analytical sensitivity of this assay. Nonetheless, these data demonstrate that phenazine production by the five Pa strains is explanatory for our spent media assay results, wherein the three inhibitory strains (191.1, 193.1, and PA14) produce phenazines at or near the MIC, and the two less-inhibitory strains (PAO1 and 145.1) produce phenazines well below the MIC.

Some studies have shown that PYO remains active under anoxic conditions [43, 49, 50], indicating that the cytotoxicity of phenazines is not solely dependent on aerobic respiration. To determine if this is the case for KPPR1, we repeated our co-culture competition assays under anaerobic conditions. Unlike previous studies, we did not observe Pa growth restriction of KPPR1 (**Figure S11A**). To confirm these studies, we repeated our spent media growth assays using the heterologous phenazine expression system. Like our co-culture results, we did not observe any growth restriction or bactericidal activity in the case of PYR under anaerobic conditions (**Figure S11B**). These results indicate that Pa restriction of Kp is dependent on environmental oxygen.

Next, we aimed to determine the translatability of these findings to a diverse set human of Pa and Kp clinical isolates derived from a variety of infection sources. First, we screened the ability of 198 clinical Kp isolates to grow in PA14 spent media. The growth of all 198 strains was highly restricted, like KPPR1 (**Figure 5A**), indicating a universal phenotype. Co-culture assays validated these findings (**Figure S12**). We then screened a subset of these clinical Kp isolates (N = 41) for growth in the spent media of our PCA, 5MPCA, and PYO constitutively expressed Ec strains. Of note, to accommodate the growth of our clinical Kp isolates, phenazine-containing spent media was generated in the absence of antibiotic selection to maintain the expression plasmids, and correspondingly, phenazine yield was lower than that in **Figure 3F**. Similar to our results with Kp 13F11, we observed that PYR and PYO significantly restricted the growth of our clinical Kp isolates (**Figure 5B**); however, this phenotype was strain-dependent, wherein some Kp strains were more resistant to PYO and PYR growth restriction than others.

**Figure 5.**
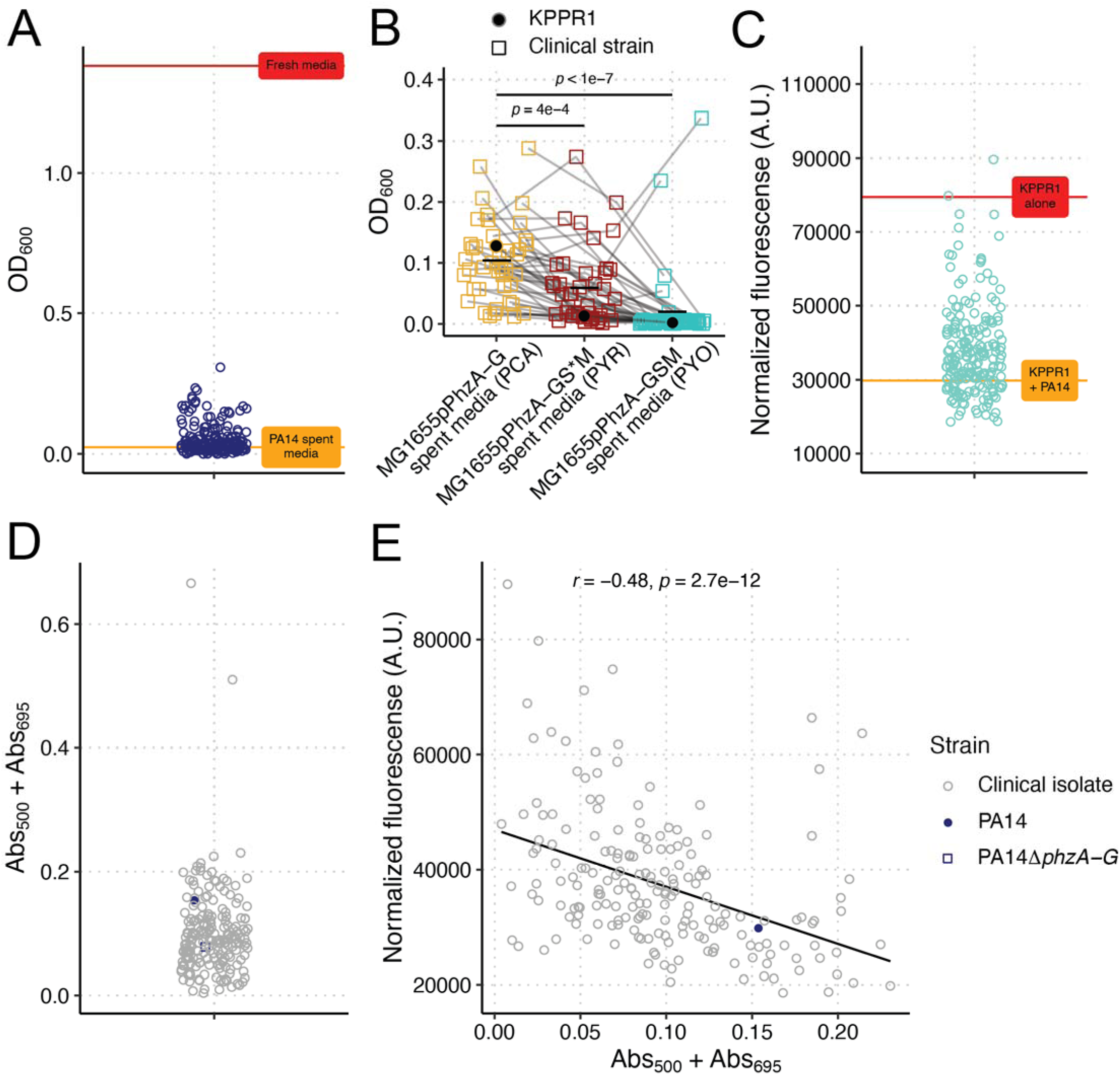
**Phenazines universally restrict Kp growth, but restriction is strain specific**. Kp clinical isolates (N = 198) were grown in sterilized spent PA14 LB (**A**). Abs_600_ was measured at 24 hours. For **A**, each data point represents the median read of each culture. The red line represents mean Abs_600_ of KPPR1 in fresh LB, and the orange line represents the mean Abs_600_ of KPPR1 in PA14 spent LB. A subset of clinical Kp isolates (N = 41) and KPPR1 were grown in filter sterilized spent media of MG1655 constitutively expressing PCA (pPhzA-G), PYR (pPhzA-GS*M), and PYO (pPhzA-GSM, **B**). Abs_600_ was measured at 24 hours. For **B**, each data point represents the median read of each culture, and grey lines connect identical isolates. *p-*values represent Tukey multiple comparison correction following one-way ANOVA. Pa clinical isolates (N = 194) were co-cultured in LB with GFP-expressing KPPR1 (**C**). Fluorescence (arbitrary units) was measured at 24 hours and normalized to culture density (Abs_600_). For **C**, each data point (blue) represents the median of each co-culture. The red line represents the mean normalized fluorescence for KPPR1, and the orange line represents the mean normalized fluorescence for KPPR1 + PA14. For Pa clinical isolates (N = 194), WT PA14, or PA14Δ*phzA-G,* phenazine production was measured in LB broth at Abs_500_ (PYR) and Abs_695_ (PYO, **D**) and correlated (N = 192) to growth restriction results from **C** (Spearman correlation test, **E**).

Then, we screened the ability of 194 Pa strains to restrict KPPR1 growth. Interestingly, KPPR1 restriction was highly strain dependent (**Figure 5C**), akin to our findings with PAO1 and PA14 (**Figure 1B-C**). Co-culture assays validated these findings (**Figure S13**). Given that phenazines are necessary for PA14 restriction of KPPR1, we then tested phenazine production in our screening conditions (**Figure S10A**). The clinical Pa strains produced phenazines to varying degrees, with many producing very little (**Figure 5D, S10B, S10D**). PYO and PYR levels as measured by absorbance were significantly correlated (**Figure S10F**). Of note, phenazine measurements in this assay are based on absorbance (**Figure S10A**), thus, there may be spectral overlap between phenazines or other non-phenazine compounds produced that are detected at the wavelengths we tested. 47 Pa strains had Abs_500_ values higher than PA14, suggesting that these strains may secrete PYR (**Figure S10D**). The ability of clinical Pa strains to restrict Kp significantly inversely correlated with phenazine production (**Figure 5E, S10C, S10E**); however, the correlation between phenazine production and Kp growth restriction was weak (*r* =-0.48, *r*^2^ =-0.23), wherein many Pa strains produce little or no phenazine but are highly restrictive and others less restrictive but appear to produce high levels of phenazine. The two strains that produced high levels of phenazines in **Figure 5D** were excluded from the analyses in **Figure 5E**, as these strains produced a pigment similar in appearance to pyomelanin, resulting in high absorbance values not due to phenazine production.

To test if other secreted Pa effectors may be a factor in Kp growth restriction as our data suggested, we selected two Pa strains that we validated as inhibitory in our co-culture assay (**Figure S13**): JV46 and JV69. In our screen, JV46 was slightly more inhibitory than PA14 (mean normalized fluorescence = 26,524.17 versus 29,785.32) and JV69 was less inhibitory than PA14 (mean normalized fluorescence = 35,396.03 versus 29,785.32), yet these strains produced similar levels of phenazines (JV46 Abs_500_ + Abs_695_ = 0.154, JV69 Abs_500_ + Abs_695_ = 0.103, PA14 Abs_500_ + Abs_695_ = 0.154). The spent media of JV69 was inhibitory to a comparative level as PA14, where JV46 was nearly completely bactericidal (**Figure S14A**). Of note, the Abs_500_ measurement of JV46 spent media was lower than that of PA14 (0.07 versus 0.086), suggesting that the bactericidal phenotype is not due to increased PYR secretion, which we confirmed by mass spectrometry (**Figure S14B**). This indicates that phenazine production is like only one mechanism by which Pa restricts Kp growth. Collectively, we concluded that at high concentrations, Kp is universally sensitive, but at low concentrations, sensitivity is strain dependent.

There is growing interest in using live biotherapeutics to preventatively decolonize Kp and other gut colonizing pathogens to lower infection risk [38, 39, 51]; however, current approaches largely depend on manipulation of the metabolic niche, rather than leveraging direct growth inhibition. Given our isolation of the wild Pa strains from mouse gut samples with low Kp colonization and their ability to directly restrict Kp, we hypothesized that these Pa strains can exclude Kp from the gut. To test this hypothesis, we colonized C57Bl6/J mice with Pa 191.1. To open the gut niche for colonization, mice were treated with ampicillin prior to Pa colonization. 191.1 was able to stably colonize the gut (**Figure S15A**), though colonization density was low. One week following Pa colonization, mice were colonized with KPPR1. An antibiotic treated group not colonized by 191.1 was included as a control. No difference in KPPR1 gut colonization density was observed between mono-colonized and co-colonized mice (**Figure S15B-C**), potentially due to low 191.1 colonization. Next, we hypothesized that the intact gut microbiome may be required to observe a restrictive phenotype, as the gut microbiome of the mice in our initial *in vivo* experiment was intact (**Figure S1A**). To this end, we used an *ex vivo* approach to test this hypothesis. Whole large intestinal contents of C57Bl6/J mice were collected and resuspended in sterile phosphate buffered saline (PBS) to generate gut microbiota-replete competition media. KPPR1 was anaerobically competed in this media against 145.1, 191.1, and 193.1. All Pa strains were viable in these conditions (**Figure S16A**); however, no reduction in KPPR1 CFUs was observed (**Figure S15D**). Together, these results indicate that the wild Pa strains are unable to restrict Kp growth in intestinal contents. Of note, this corroborates our finding that phenazine-dependent Kp growth restriction is dependent on environmental oxygen.

Given our gut findings that were contradictory to our original observations, we turned to other relevant body sites for interrogating the interactions between Pa and Kp where phenazine production may be important. Interestingly, stratification of clinical Pa growth restriction data based on the clinical site of origin suggested site specificity (**Figure 6A**, T**able S1**). Pa isolated from the blood and urine were more restrictive compared to those isolated from the respiratory tract or wounds. To determine if there is a causal relationship between phenazine production and site-specific restriction of Kp growth, we aimed to test phenazine-dependent Kp growth restriction in conditions that mimic respiratory and urine infections. To this end, co-culture competitions were performed in murine bronchoalveolar lavage fluid (BALF) and *ex vivo* bladder homogenate. Both *ex vivo* tissues sustained the growth of both Kp and Pa (**Figure S16B-C**). Interestingly, we did not detect phenazine-dependent growth restriction of Kp in murine BALF (**Figure 6B**), supporting a diminished role in Kp lung co-infection. Conversely, we detected a significant phenazine-dependent growth restriction phenotype in *ex vivo* bladder homogenate (**Figure 6C**), consistent with a more important role of phenazines during urinary tract infection. Collectively, these data demonstrate that the impact of phenazines on Pa-Kp interaction, and therein, infection outcomes, is site specific.

**Figure 6.**
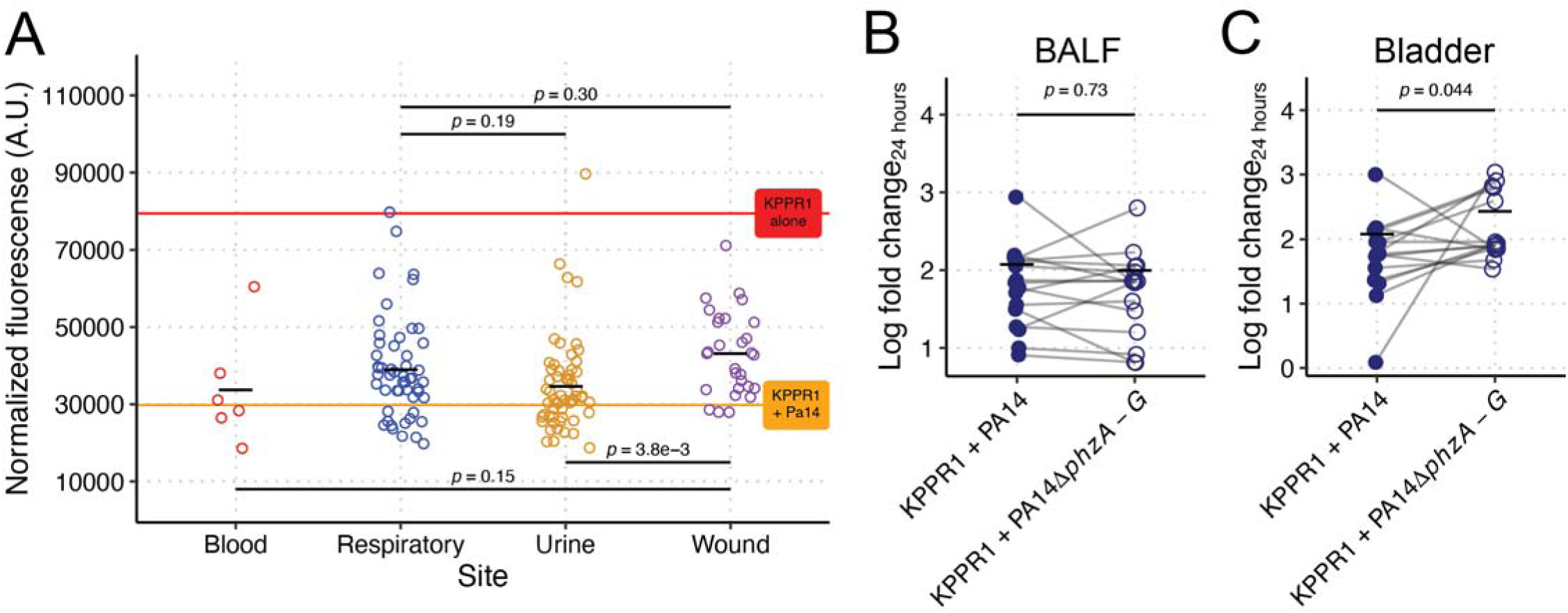
Phenazine-dependent restriction of Kp is site-specific. Mean normalized fluorescence for KPPR1 from co-culture with Pa clinical isolates (N = 194) in LB from **Figure 5C** was stratified by clinical culture site (**A**). The red line represents mean normalized fluorescence for KPPR1, and the orange line represents the mean normalized fluorescence for KPPR1 + PA14. KPPR1 was grown in co-culture in bronchoalveolar lavage fluid (**B**) or *ex vivo* bladder homogenate (**C**) with WT PA14 or PA14Δ*phzA-G*. For **B-C**, “Log fold change_24 hours_” = log_10_(output KPPR1 CFU at 24 hours/input KPPR1 CFU). Each data point is a biological replicate, connecting lines indicate corresponding *ex vivo* samples, and horizontal lines indicate the mean of each dataset. *p-*values represent unpaired *t* tests.

## Discussion

In this study, we investigated the ability of Pa to restrict Kp growth and demonstrated that it can do so in a contact-independent manner. These findings are partially dependent on nutrient competition; however, the synergistic and/or additive effects of Pa-secreted phenazines account for the majority of this growth restriction. The phenazine PYO, well known for its antibacterial activity, has bacteriostatic effects on Kp, while the less well-characterized PYR has bactericidal effects, at least at high concentrations. Interestingly, these phenotypes are highly dependent on the Pa strain, wherein phenazine production is a strong correlate of Kp restriction, though again, is not completely explanatory. Rather, some Pa strains restrict Kp growth without producing high levels of phenazines. Further complicating this relationship, environmental conditions also impact the outcome of Pa-Kp competition. We observed phenazine-dependent growth restriction in conditions that mimic bladder infection but not lung infection, which corresponded to experimental interrogation of our collection of clinical isolates. Collectively, our study shows that interactions between Pa and Kp are highly complicated, and that variables like genetic composition and environmental conditions are key determinants of fitness outcomes. This provides a potential explanation for study-to-study variation and lays a foundational approach for interrogation of microbial interactions during pathogenesis wherein many variables that may impact experimental outcomes should be considered.

The toxicity of PYO to various bacterial species has been shown previously, whilst 5MPCA toxicity has been previously observed in *Candida albicans* [42, 47]. It has been suggested that in Pa, 5MPCA is rapidly converted to PYO to prevent accumulation of a toxic intermediate [52]. The spontaneous frameshift in PhzS in our MG1655pPhzA-GS*M strain leads to increased production of 5MPCA and increased Kp toxicity, supporting the hypothesis that it is more toxic than PYO. PYO has been shown to accept electrons from the electron transport chain and donate them to oxygen, disrupting cellular redox balance and generating cytotoxic reactive oxygen species [53]. Whilst PYR toxicity was dependent on aerobic conditions, its high redox potential [54] make oxygen reduction unfavorable [55]. Furthermore, Ec MG1655 produced large titers of PYR that were bactericidal to Kp, despite genetic and metabolic similarities between these species [38]. This suggests a specific mode of PYR toxicity other than reactive oxygen species generation. One major difference between aerobic respiration in Ec and *Klebsiella* is the dependence of the latter on type II NADH-dehydrogenases, especially in urinary media [56]; however, these studies were performed with *K. aerogenes*. In Pa, these enzymes are the predominant phenazine reductases, suggesting phenazine inhibition of Kp could be caused by it acting as an electron acceptor, uncoupling NADH oxidation to ATP synthesis in the electron transport chain [57]. Further work is required to determine the mechanism underlying Kp phenazine toxicity.

The initial observation that was the inception of this study was the isolation of the wild Pa strains (145.1, 191.1, 193.1) from murine large intestinal contents (**Figure S1A**). Yet, deeper interrogation revealed that phenazine-dependent growth restriction of Pa by Kp is oxygen-dependent (**Figure S11**). This result is surprising, as our original Kp gut colonization experiments were performed in the context of an unperturbed gut microbiome, which would maintain a largely anoxic environment. Our original goal was to test if gut colonization by Pa would exclude Kp from the gut, with a broader goal to evaluate the underpinning mechanisms as potential decolonization strategies. Yet, we found that Pa is not readily adaptable to our gut colonization model, which was confirmed in discussion with other groups that had attempted Pa gut colonization. Of note, we were unable to exactly replicate the experiments in **Figure S1A** with our wild Pa strains, as the source vendor of these mice no longer breeds C57Bl6 mice in the barrier (room of origin) from which the original mice were sourced. As the barrier of origin is a known confounder that impacts gut microbial community structure [58], it is challenging to specifically determine if our original observations are repeatable. However, our attempts to recapitulate these findings suggest that our original observation was a “True, true, unrelated” situation, wherein Pa could have been a contaminant or transient gut colonizer, inhibiting Kp through a phenazine-independent modality, and/or another microbe or set of microbes was explanatory for the reduction in colonization that we observed in our original experiments. These accidental observations led to interesting findings, nonetheless.

Despite not identifying a microbial interaction that led to a reduction in Kp gut colonization, there are significant implications of these data for our understanding of how microbial interactions shape infection outcomes. In the lung, a growing number of studies recognize the importance of the lung microbial community in health outcomes, where it is associated with lung transplant failure, lung cancer, asthma, idiopathic pulmonary fibrosis, and infectious pneumonia, amongst others (reviewed in [2]). This concept is best studied in cystic fibrosis (CF). CF disease severity, as well as the immunological and treatment response, are all strongly influenced by microbial interactions between important pathogens, such as Pa, *Staphylococcus aureus*, and others (reviewed in [59, 60]). During chronic Pa lung infection in CF patients, Pa adapts to a more cooperative state, resulting in reduction of phenazine expression [61–63]. In humans, lung infections appear to be exacerbated by Pa-Kp co-infection [11–13]. In a mouse model of lung co-infection, Pa exacerbated Kp infection, but the bacterial mechanisms underpinning that exacerbation are unknown [17]. In the urinary tract, it is well known that microbial interactions greatly shape infection outcomes, including disease severity and secondary effects of infection, such as formation of struvite crystals (reviewed in [64]). For example, *Providencia stuartii* enhances *Proteus mirabilis* dissemination during UTI, whereas *Morganella morganii* reduces *P. mirabilis* dissemination during UTI [65, 66]. Kp and Pa urinary tract co-infections are known to occur [14], but little is known about what shapes their interactions and their infectious outcomes. Our data suggest there may be important impacts of phenazines on the local polymicrobial microenvironment during infection that have yet to be tested using these models; however, this requires further investigation. Interestingly, PYO is necessary for fitness during Pa lung infection [67]; thus, PYO (or other phenazines) may underpin the exacerbation of Kp infection by Pa. Conversely, RhlR (and therein, phenazines) is dispensable for fitness in a catheter-associate UTI mouse infection model using Pa14 [68]. Thus, it may be that phenazines are important for shaping polymicrobial UTI outcomes, rather than driving fitness during mono-microbial infection. Future studies should aim to disentangle the complex interactions between phenazines, the host, and other microbes during polymicrobial infection when Pa is present.

In other organ systems, microbial interactions have become key targets for health-oriented interventions. The gut is the most prominent focus of such efforts, including the successful use of fecal microbiota transplantation to treat recurrent *Clostridioides difficile* infection and ongoing clinical trials of live bacterial therapeutics (reviewed in [69]). Such interventions are at the nascent stage of consideration for both Kp and Pa infections, and microbial interactions are being considered as anti-infection therapeutic targets (reviewed in [70], for example). Our study shows that it may be necessary to consider the type of infection and strain genetic repertoire when evaluating the efficacy of curative or anti-infection therapies, as efficacy is likely site and strain specific. Additionally, our use of a heterologous overexpression system indicates that future studies should determine if engineered Ec or bacterial species with increased antimicrobial secondary metabolite production, such as PYO or PYR, could be used to eliminate Kp or other microbes in human-relevant systems.

There is an urgent need for new approaches to prevent deaths from antibiotic resistant bacterial infections as global antimicrobial resistance rises [71]. Both Kp and Pa are important causes of antibiotic-resistant infection, both ranking in the top six most prevalent causes of deaths attributable to antibiotic resistance [19, 20]. There are likely opportunities to shorten, prevent, or cure infections with these antibiotic-resistant pathogens through targeted manipulation of polymicrobial competitions that underpin and/or enhance their virulence. A major barrier in therapeutic leveraging of microbial competition is the lack of mechanistic studies of these interactions, which are necessary to understand and predict their efficacy. Additionally, a significant challenge for interventions that rely on microbial competition is the tremendous patient heterogeneity due to variations in genetics, immunological baseline, behavior, and microbial ecosystems (reviewed in [72]). As such, it is important to measure the effects of these variables when evaluating the efficacy of interventions that leverage microbial competition, when possible. If successful, such interventions have the potential to both lower the burden of infection complicated by antibiotic resistance and re-potentiate existing antibiotic therapies by reducing their use.

## Materials and methods

### Ethics statement

This study was performed in strict accordance with the recommendations in the *Guide for the Care and Use of Laboratory Animals* [73]. The Indiana University Institutional Animal Care and Use Committee approved this research (protocol #22114, PI: Jay Vornhagen). Wild Pa strains (145.1, 191.1, 193.1) were isolated from mouse experiments (**Figure S1A**) approved by The University of Michigan Institutional Animal Care and Use Committee (protocol #PRO00007474, PI: Michael A. Bachman). Clinical isolates were collected and de-identified by RFR in accordance with approval by the Indiana University Institutional Review Board (protocol #16139, PI: Ryan F. Relich).

### Materials, media, and bacterial strains

All chemicals were purchased from Sigma-Aldrich (St. Louis, MO) or Fisher Scientific (Fairlawn, NJ) unless otherwise indicated. Bacterial strains used in this study are described in **Table 2**. Bacteria were cultured in Luria-Bertani (LB) broth, or in M9 minimal medium (M9 salts, 0.2 M MgSO_4_, 0.01 M CaCl_2_, with 1% casamino acids, “M9-cas”) at 37° C with shaking at 220 rpm, or on LB agar at 27° C (Kp) or 37° C (Pa and Ec) supplemented with kanamycin (40 or 25 µg/ml) and/or rifampicin (30 µg/ml) as appropriate. pJL1-sfGFP was a gift from Michael Jewett (Addgene plasmid 102634). pSJ102 containing Ec was grown at 30° C for optimal phenazine production. For experiments that required species specific selection, KPPR1 was selected on LB agar with 30 µg/ml rifampicin grown at 27° C and Pa was selected on Pseudomonas Isolation Agar (Becton, Dickinson and Company, Franklin Lakes, NJ).

**Table 2.** Bacterial strains used in this study.

| **JV Number** | **Species** | **Strain** | **Source or reference** |
| --- | --- | --- | --- |
| JV1 | *Klebsiella pneumoniae* | KPPR1 | [76] |
| JV3 | *Pseudomonas aeruginosa* | 145.1 | This study |
| JV4 | *Pseudomonas aeruginosa* | 191.1 | This study |
| JV5 | *Pseudomonas aeruginosa* | 193.1 | This study |
| JV10 | *Klebsiella pneumoniae* | 13F11 | [77] |
| JV11 | *Escherichia coli* | MG1655 | [78] |
| JV13 | *Pseudomonas aeruginosa* | PAO1 | [79] |
| JV14 | *Pseudomonas aeruginosa* | PA14 (used only for JV388 and JV389 comparisons) | Waters Lab, Michigan State University |
| JV338 | *Pseudomonas aeruginosa* | PA14Δ*rhIR* | [45] |
| JV339 | *Pseudomonas aeruginosa* | PA14Δ*rhII* | [45] |
| JV450 | *Pseudomonas aeruginosa* | PA14 (SMC232) | [80] |
| JV449 | *Escherichia coli* | DH5apJL1-sfGFP | [81] |
| JV461 | *Klebsiella pneumoniae* | KPPR1pJL1-sfGFP | This study |
| JV462 | *Pseudomonas aeruginosa* | PA14Δ*phzA1*-G1Δ*phzA2*-G2 ("PA14Δ*phzA*-G," SMC5020) | [82] |
| JV463 | *Pseudomonas aeruginosa* | PA14 SXO *phzS* ("PA14Δ*phzS*," SMC5123) | [83] |
| JV466 | *Pseudomonas aeruginosa* | PA14Δ*phzH*Δ*phzM* SXO *phzS* (SMC5126) | [83] |
| JV467 | *Pseudomonas aeruginosa* | PA14Δ*phzH* (SMC5127) | [83] |
| JV468 | *Pseudomonas aeruginosa* | PA14Δ*phzM* (SMC5128) | [83] |
| JV531 | *Escherichia coli* | SJ102 (MG1655 intC::λpR-YFP-Cmr) + pSB3K3_EVC ("pEmpty") | This study |
| JV534 | *Escherichia coli* | SJ102 + pSB3K3_*phzA*-GSM ("pPhzA-G") | [74] |
| JV537 | *Escherichia coli* | SJ102 + pSB3K3_*phzA*-GS*M ("pPhzA-GS*M") | This study |
| JV540 | *Escherichia coli* | SJ102 + pSB3K3_*phzA*-GSM ("pPhzA-GSM") | [74] |

### Phenazine synthesis plasmids

Plasmids for PCA (pSB3K3_*phzA-G*) and PYO (pSB3K3_*phzA-GSM*) expression were obtained from a previous study [74]. The empty vector control (pSB3K3_EVC) was made by performing a NotI digestion of pSB3K3_*phzA-GSM*, followed by ligation with a neutral oligonucleotide linker made by annealing the following primers: 5’-GGCCGCGTACGACTTACGC-3’ and 5’-GGCCGCGTAAGTCGTACGC-3’. All plasmids were transformed into *E. coli* SJ102 (MG1655 intC::λpR-YFP-Cmr [75]) by electroporation. The PYR expression plasmid (pSB3K3_*phzA-GS*M*) was obtained spontaneously by selecting a colony of pSB3K3_*phzA-GSM* cells which displayed a red corona on LB-agar plates. Plasmid sequences were confirmed by full plasmid sequencing.

### Whole genome sequencing

Pa strains were cultured overnight in LB broth for 24 hours prior to DNA extraction. DNA was extracted using Monarch Genomic DNA Purification Kit (New England Biolabs, Ipswich, MA) according to the manufacturer’s instructions with the only deviation being the substitution of 60 °C elution buffer for 34 °C DNase/RNase-Free Water (Zymo Research, Irvine, CA). The extracted DNA quantity was analyzed using a Quibit 4 Fluorometer 1x dsDNA HS analysis. DNA extractions above a concentration of 8 ng/uL were prepared for sequencing with a Rapid Barcoding Kit 24 V14 (Oxford Nanopore Technologies, Oxford, England) utilizing their Rapid sequencing DNA V14 – barcoding protocol (RBK_9176_v114_revQ_27Dec2024). Bacterial genomes were sequenced with a Flo-Min114 flow cell on a MinION Mk1B device (Oxford Nanopore Technologies, Oxford, England) and base called using the ONT MinKNOW software v24.06.5 (Oxford Nanopore Technologies, Oxford, England). The reads were assembled into scaffolds using Flye Assembler v2.9.5 [84].

### Phylogenetic analysis

We determined that the in-house assembled genomes were most similar to ST175 by multi-locus sequence typing [85]. These assembled genomes were combined with genomic sequences of five ST175 Pa strains downloaded from the bacterial and viral bioinformatics resource center on December 9, 2023 [86]. The genomic sequences of an additional 65 Pa strains were downloaded from the National Center for Biotechnologies Information on December 6, 2023 [87]. These 73 genomes, in addition to the three in-house assembled genomes, were annotated using Prokka v1.14.5 [88] and the core genome determined using Roary v3.13.0 [89]. The core genome in this study was defined as loci with 90% consistency between isolates. A maximum likelihood phylogenetic tree was assembled using FastTree v2.1.11 and visualized using iTOL v7.0 [90, 91].

### Interspecies competition assays

For co-culture competition assays, strains were cultured overnight LB or M9-cas. Strains were then co-inoculated 1:1,000 in 1 mL of the medium of interest and mixed. This input was serially diluted and selectively spot-plated for CFU quantification. After 24 hours of shaking incubation, the output was again serially diluted and plated. For spent-media competition assays, strains were cultured overnight in the medium of interest. Spent media was created by filtering the supernatant of centrifuged culture with a 0.22 μm syringe filter. The strain of interest was inoculated 1:1,000 into spent media and was then plated and incubated as described above. For MG1655pPhzA-GS*M spent media assays, except those in **Figure 3F**, dithiothreitol was added to final concentration of 5 mM.

### Growth curves

Strains were cultured overnight in culture medium of interest (LB or M9-cas) then diluted to an OD_600_ of 0.01 in the medium of interest the following day. Cultures were incubated at 37° C with aeration and Abs_600_ readings were taken every 15 min using a BioTek microplate reader with Gen5 software (Version 3.12.08, BioTek, Winooski, VT) for 24 hours. To simultaneously assess doubling time, growth rate, lag time, non-sigmoidal growth due to stress and bacterial density, area under the curve analysis was used to quantify differences in growth, as in [33].

### BioLog Phenotype MicroArray analysis

BioLog Phenotype MicroArrays (Biolog, Hayward, CA) were performed according to manufacturer’s instructions with some modifications, as in [33]. KPPR1, PAO1, PA14, and the mouse-derived Pa strains (Pa 145.1, Pa 191.1, and Pa 193.1), were cultured overnight in LB, then bacteria were pelleted, washed once in sterile PBS, and re-suspended in sterile PBS to avoid aberrant transfer of xenonutrients. Each strain was diluted in IF-0 medium to a final OD_600_ of 0.035 and 100 μL was plated onto plates PM1 and PM2 with gentle mixing. After inoculation, plates were statically incubated overnight at 37° C. Following 24 hours of incubation, growth was measured at OD_595_.

### PA14NR library screen

Construction of the PA14NR transposon library used in this study has been extensively described elsewhere [92]. Library plates were pin-replicated onto black 96-well microplates containing 100 of µL LB agar. After 24 hours of static growth at 37° C, liquid culture of KPPR1pPJL1-sfGFP was diluted 1:1,000 in LB broth and 100 µL was added on top of the agar. After 24 hours of static 37° C incubation, a fluorescence read was performed (Excitation: 479, Emission: 520, Optics: Bottom, Gain: 50).

### High-resolution mass spectrometry

For high-resolution mass spectrometry (HR-MS) detection of phenazines in cell-free supernatant, bacteria were grown overnight in LB broth at 30° or 37° C with cognate antibiotics. Cultures were centrifuged at 7,000 x*g* for 5 minutes and supernatant was filtered through a 0.22 μm syringe filter. This filtrate was then centrifuged at 16,000 x *g* for 10 minutes to remove debris, and 500 μL was diluted 1:1 with LC-grade methanol. Sample (1 μL) was injected onto 1290 LC - 6545 QTOF. The ESI-HRMS analyses were performed in positive ion mode, utilizing nitrogen as the nebulizing and drying gas. The instrumental conditions were as follows: nebulizer pressure, 25 psi; drying temperature, 325° C; drying gas, 8 L/min; sheath gas temperature, 400° C; sheath gas flow, 12 L/min; fragmentor, 220 V; skimmer, 65 V; Oct 1 RF Vpp, 750 V. For the first HPLC condition, an Agilent ZORBAX Eclipse Plus C18 column (HD 2.1*5 0mm 1.8-Micron) was used for separation. 0.1% formic acid in water (v:v) as mobile phase A and 0.1% formic acid acetonitrile as mobile phase B (v:v) were used. The flow rate is 0.6 mL/min. The gradient starts as 5% B and holds for 0.5 minute, increased from 5% B to 95 %B in 4.5 minutes and holds for 0.5-minute, 95%B to 5%B in 0.5 minute and holds for 0.5 minute. For the second HPLC condition and ACQUITY UPLC BEH Amide column (150 mm × 2.1, 1.7 μm) was used for separation. The mobile phases were heated to 45 degrees. 0.1% formic acid in water (v:v) as mobile phase A and 0.1% formic acid acetonitrile as mobile phase B (v:v) were used. The flow rate is 0.6 mL/min. The gradient starts as 5% A and holds for 1 minute, increased from 5% A to 95 % A in 9 minutes and holds for 1 minute, 95% A to 5% A in 1 minute and holds for 1 minute. Data were analyzed using mzmine [93]. Extracted ion chromatographs were generated for PCA ([M+H+] = 225.0586 m/z), 5MPCA ([M+] = 239.0815 m/z), and PYO ([M+] = 211.0866 m/z) with 10 ppm error.

### MIC and MBC assays

MIC assays were performed as previously described [94]. For PYO, pure PYO (5-methyl-1(5H)-phenazinone, Cayman Chemical, Ann Arbor, Michigan) was resuspended to a concentration of 16 mg/mL in dimethylsulfoxide and then diluted into LB or PA14Δ*phzA-G* spent media to 1 mg/mL. 100 μL of this solution was plated into U bottom 96-well plate in triplicate and then 2-fold serially diluted into LB or PA14Δ*phzA-G* spent media 10 times. Then, 50 μL of KPPR1 culture inoculated in LB or PA14Δ*phzA-G* spent media to final concentration of OD_600_ = 0.02 was plated across each serial dilution, yielding a final OD_600_ = 0.01, and incubated at 37° C shaking for 24 hours. After 24 hours, MIC was determined to be lowest concentration where growth was not visually observed.

For PYO MBC assays, the entire MIC assay described above was spot plated onto LB agar at 24 hours and grown overnight at 37° C. After overnight growth, MBC was determined to be the lowest concentration at which no Kp colonies were recovered.

For PYR, 100 μL of 100% MG1655pPhzA-GS*M spent media was plated into U bottom 96-well plate and serially diluted in MG1655pEmpty spent media with 10 mM DTT 7 times. Then, 50 μL of KPPR1 culture inoculated in MG1655pEmpty spent media to final concentration of Abs_600_ = 0.02 was plated across each serial dilution, yielding a final DTT concentration of 5 mM and Abs_600_ = 0.01. Additionally, KPPR1 culture was inoculated in 100 μL of 100% MG1655pPhzA-GS*M spent media with 5 mM DTT to final OD_600_ = 0.01 to ensure a 100% MG1655pPhzA-GS*M spent media condition. All media were incubated at 37° C shaking for 24 hours. After 24 hours, MIC was determined to be lowest concentration where growth was not visually observed.

To assess additive and/or synergistic effects of PYO and PYR, a checkboard assay was used. To this end, a 2-fold serial dilution of PYO, starting at 62.5 μg/mL was combined with 2-fold serial of MG1655pPhzA-GS*M spent media, starting at 100%. DTT was added to a final concentration of 5 mM and KPPR1 culture was inoculated to final OD_600_ = 0.01. This array was incubated at 37° C shaking for 24 hours. After 24 hours, MIC was determined to be lowest concentration where growth was not visually observed.

### Phenazine quantification

The absorbance spectra of pure PYO and MG1655pPhzA-GS*M spent media was measured in both reducing (5 mM DTT) and oxidizing conditions (0.3% peroxide). We determined that oxidized MG1655pPhzA-GS*M spent media and untreated PYO yielded the greatest differences between these spectra at Abs_500_ and Abs_695_, respectively. For PYO quantification, Pa cells were removed from spent Pa media generated under conditions identical to spent media assays. The raw Abs_695_ of spent media was measured or compared to a standard curve of pure PYO to estimate PYO concentrations. For 5MPCA quantification, Pa spent media was oxidized, then the raw Abs_500_ of spent media was measured or compared to a standard curve of oxidized MG1655pPhzA-GS*M spent media to interpolate estimate 5MPCA concentrations.

### Collection and screening of clinical isolates

The IU Health Division of Clinical Microbiology collected clinical isolates between May and July 2023 from IU Health facilities across the state of Indiana. Of these, 198 isolates were identified by MALDI-TOF (Bruker) as Kp, and 194 as Pa. These isolates were derived from a variety of infection sites and patient information was de-identified. Isolates were passaged once from primary plating media (sheep blood agar or MacConkey agar) prior to storage on brain-heart infusion agar slants (ThermoFisher) at room temperature.

To screen Kp growth in PA14 spent media, clinical Kp isolates and KPPR1 were cultured overnight in LB and inoculated 1:100 into sterile spent PA14 media or fresh LB. After 24 hours of shaking incubation at 37° C, Kp growth was measured at OD_600_.

To screen Kp growth against specific phenazines, a subset of 49 clinical Kp isolates and KPPR1 were cultured in sterile spent MG1655pEmpty media. Eight isolates were determined to be unable to grow in this spent media and excluded from future assays. The remaining 41 clinical Kp isolates were cultured in sterile spent MG1655pPhzA-G, MG1655pPhzA-GS*M, or MG1655pPhzA-GSM media generated in the absence of antibiotic selection for their respective expression plasmids to permit clinical isolate growth. After 24 hours of shaking incubation at 37° C, Kp growth was measured at Abs_600_.

To screen clinical Pa isolate restriction of KPPR1 growth, clinical Pa isolates and KPPR1pJL1-sfGFP were cultured overnight in LB and inoculated 1:1,000 in LB for a 1:1 competition of KPPR1pJL1-sfGFP in co-culture with one clinical isolate. After 24 hours of shaking incubation at 37° C, an OD_600_ reading was taken, as was a fluorescence read (Excitation: 479, Emission: 520, Optics: Top, Gain: 50).

### *Ex vivo* interspecies competition assays

Strains for *ex vivo* interspecies competition assays were cultured overnight in LB, washed in sterile PBS and re-suspended in sterile PBS before inoculation to avoid aberrant transfer of xenonutrients.

*Large intestinal contents:* 6-to-8-week-old C57Bl6/J male mice were sourced from the Jackson Labs from barrier RB07. Following acclimation, mice were euthanized, the large intestine (cecum and colon) was removed, and its contents were gently homogenized in 2 mL sterile, pre-reduced PBS and divided into 400 μL aliquots. Immediately following collection, samples were transferred to an anerobic chamber and inoculated to a final concentration of ∼5×10^7^ CFU KPPR1 alone or an equal ratio of KPPR1 with each wild Pa strain. Samples were incubated at 37° C for 48 hours, then dilution plated on species selective media to quantify final bacterial densities. Only male mice were used to account for sex-derived microbiome differences.

*BALF:* 6-to-8-week-old C57Bl6/J male mice from the Jackson Labs’ barrier RB07 and C57Bl6 male mice from Charles River from barrier K61 were sourced for BALF collection. Two vendor-barrier sources were used to determine if there were vendor-specific effects of the lung microbiome, as has been previously reported [95]. Following acclimation, mice were euthanized, and BALF was collected as previously described [33]. If BALF recovery was > 1 mL, sterile PBS was added to 1 mL. BALF was divided into 400 μL aliquots and inoculated 1:1,000 with a 1:1 mix of KPPR1 and WT PA14 or PA14Δ*phzA-G*. Input bacterial densities were determined by dilution plating on species selective media. Samples were incubated aerobically with agitation (220 rpm) at 37° C for 24 hours, then dilution plated on species selective media to quantify final bacterial densities. Only male mice were used to account for sex-derived microbiome differences, and no vendor-based differences were detected.

*Bladder homogenate:* Bladders were collected from the mice used for BALF collection. Bladders were collected into 1 mL sterile PBS, then thoroughly homogenized. Homogenized tissue was centrifuged at 21,300 x *g* and resulting supernatant was collected. If homogenate recovery was 1 mL, sterile PBS was added to 1 mL. Bladder homogenate was divided into 400 μL aliquots and inoculated 1:1,000 with a 1:1 mix of KPPR1 and WT PA14 or PA14Δ*phzA-G*. Input bacterial densities were determined by dilution plating on species selective media. Samples were incubated aerobically with agitation (220 rpm) at 37° C for 24 hours, then dilution plated on species selective media to quantify final bacterial densities. As above, no vendor-based differences were detected.

### In vivo models

*Gut colonization model:* Gut colonization was performed as previously described [33, 58], with some minor modifications. Briefly, 6-to-8-week-old C57Bl6/J (equal numbers of male and female) mice were sourced from the Jackson Labs from barrier RB07. Following acclimation, prior to colonization, mice were administered 0.5 g/L ampicillin via drinking water. Four days following antibiotic administration, mice were orally gavaged with ∼10^7^ CFU stationary-phase Pa 191.1 suspended in 250 μL sterile PBS. Fecal pellets were collected at predetermined timepoints, homogenized in 300 μL sterile PBS, and Pa gut density was measured via dilution plating on selective medium. Seven days after 191.1 administration, mice were orally gavaged with ∼10^8^ CFU stationary-phase KPPR1 suspended in 250 μL sterile PBS. Fecal pellets were collected at predetermined timepoints, homogenized in 300 μL sterile PBS, and Kp gut density was measured via dilution plating on selective medium. Mice were euthanized seven days after KPPR1 administration, ceca were collected and homogenized in 1 mL sterile PBS, and Kp density was measured via dilution plating on selective medium.

## Statistical analysis and data availability

All *in vitro* experimental replicates represent biological replicates. For *in vitro* studies two-tailed Student’s *t* test or ANOVA followed by Tukey’s multiple comparisons post-hoc test on log_10_ transformed data was used to determine significant differences between groups. For *ex vivo* and *in vivo* studies, all experiments were replicated at least twice, accounting for sex as a biological variable when appropriate. A *p-*value of less than 0.05 was considered statistically significant for the above experiments, and analysis was performed using base R version 4.5.0. Data were processed, analyzed, and visualized using R packages “readxl,” “tidyverse,” “dplyr,” “vegan,” “outliers,” “EnvStats,” “ggplot2,” “ggrepel,” and “ggtext.” All raw data and analysis scripts used for this study are available at https://github.com/jayvorn/Pseudomonas-aeruginosa-and-Klebsiella-pneumoniae-phenazines (data embargoed until September 1, 2026). Pa assemblies are available on the Sequence Read Archive (Bioproject PRJNA1311040 [embargoed until September 1, 2026]).

## Acknowledgements

The authors would like to acknowledge Dr. Natasha Tilston and the members of her lab for their thoughtful insight into this project even though they are virologists. The authors also acknowledge Dr. Kelly Bachta for sharing their protocols and insight into *in vivo P. aeruginosa* experiments. The authors thank Professor Christopher J. Howe and Dr. Robert W Bradley for providing phenazine production plasmids. Finally, the authors acknowledge the Indiana University Pervasive Technology Institute for providing supercomputing resources that have contributed to the research results reported within this paper.

## Contributions

Conceptualization: JV, CW, DHL

Methodology: JV, LZ, CW, DHL

Investigation: KT, OS, JML, BW, VVC, JA, MN, RFR, LZ, CW, JV

Visualization: CW, JV

Funding acquisition: JV

Project administration: JV

Supervision: JV

Writing – original draft: KT, OS, CW, JV

Writing – review & editing: All authors

## Use of AI

ChatGPT-4o was used to edit the R scripts associated with this manuscript to enhance their accessibility following human-driven drafting and analysis. Functionality of all edited scripts was confirmed prior to publication.

