## Supplemental data for "*Pseudomonas aeruginosa* phenazines dictate site-specific competitive interactions with *Klebsiella pneumoniae*"

Supplementary Information

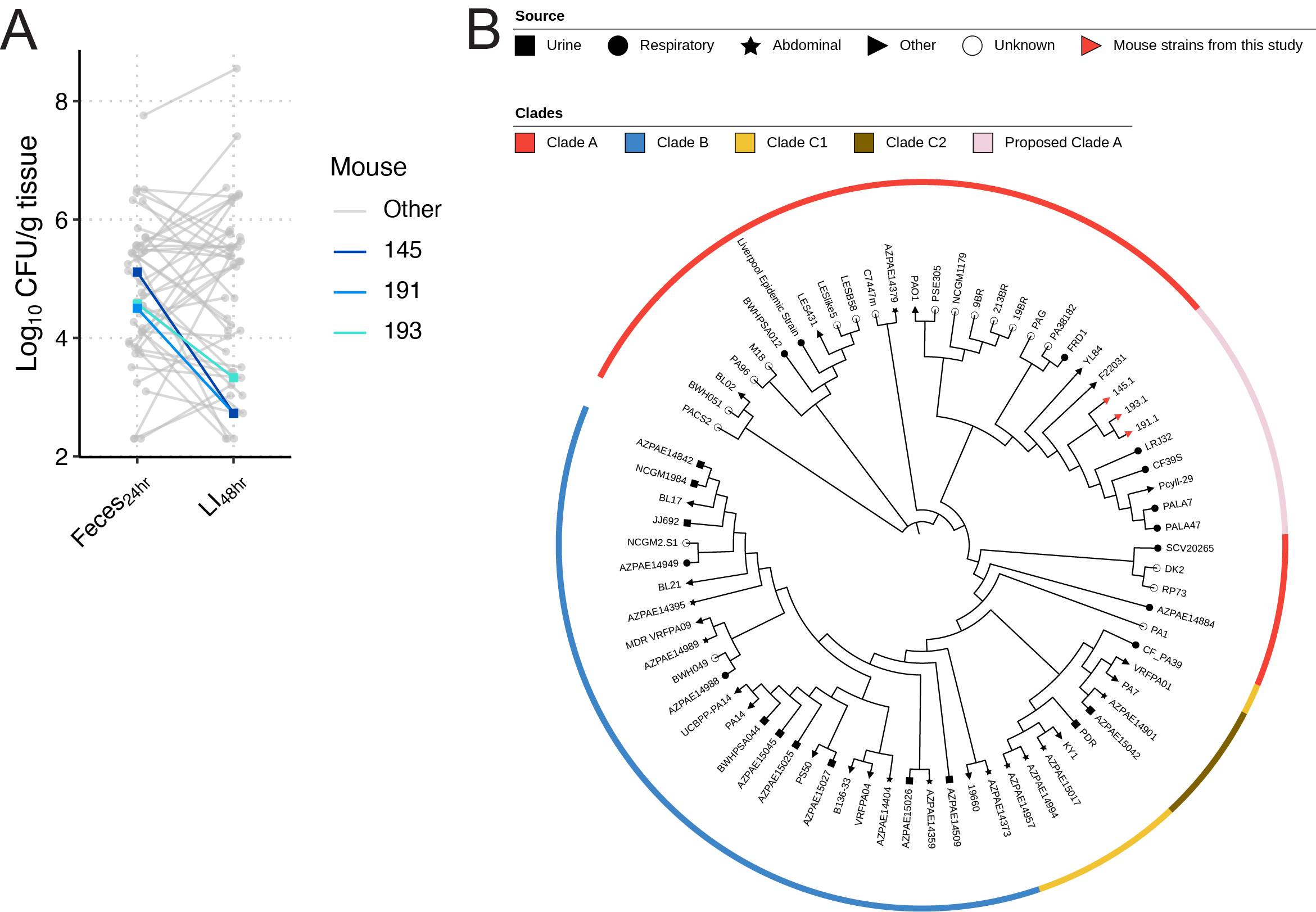

**Figure S1. Detection and phylogenetic characterization of wild Pa strains in Kp-colonized mice.** 6-to-8-week-old C57 mice from Taconic Farms were orally inoculated with 10^8^ CFU WT KPPR1 or 13F11. Kp was enumerated from feces at 24 hours and large intestinal contents (LI) 48 hours post-inoculation (**A**). Pa was isolated from mouse 145, 191, and 193 and subjected to whole genome sequencing. These strains and subset of Pa isolates from a previous study (31173069) were used to build an approximately-maximum-likelihood phylogenetic tree based on a core genome alignment of these strains to determine if strains 145.1, 191.1, and 193.1 group with Clade A (PAO1 clade), Clade B (PA14 clade), or Clade C (split Pa7 clade). Select Pa strains absent in the previous study that represent ST175 were also included, as strains 145.1, 191.1, and 193.1 were predicted to be most like ST175 using multi-locus sequence typing. The ST175 representative and mouse strains grouped with Clade A, thus they are labelled “Proposed Clade A” (**B**).

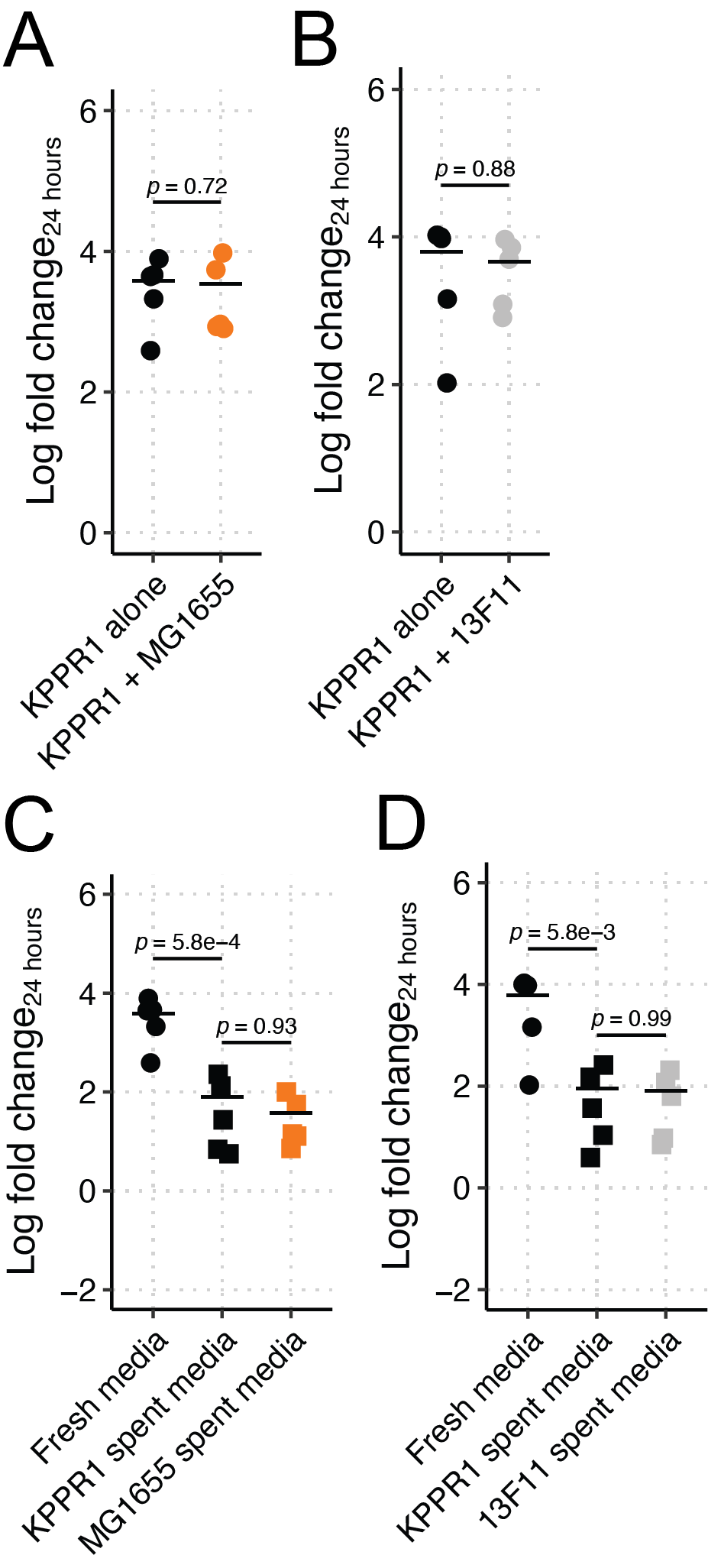

**Figure S2. Ec MG1655 and an isogenic neutral Tn mutant do not restrict KPPR1 growth in LB.** Kp KPPR1 was grown alone or in co-culture in LB with Ec MG1655 (**A**), Kp 13F11 (**B**), or in filter sterilized spent media of KPPR1, Ec MG1655 (**C**), or Kp 13F11 strain (**D**)**.** For **A-D**, “Log fold change_24 hours_” = log_10_(output KPPR1 CFU at 24 hours/input KPPR1 CFU). *p-*values represent Tukey multiple comparison correction following one-way ANOVA. Each data point is a biological replicate, and horizontal lines indicate the mean of each dataset.

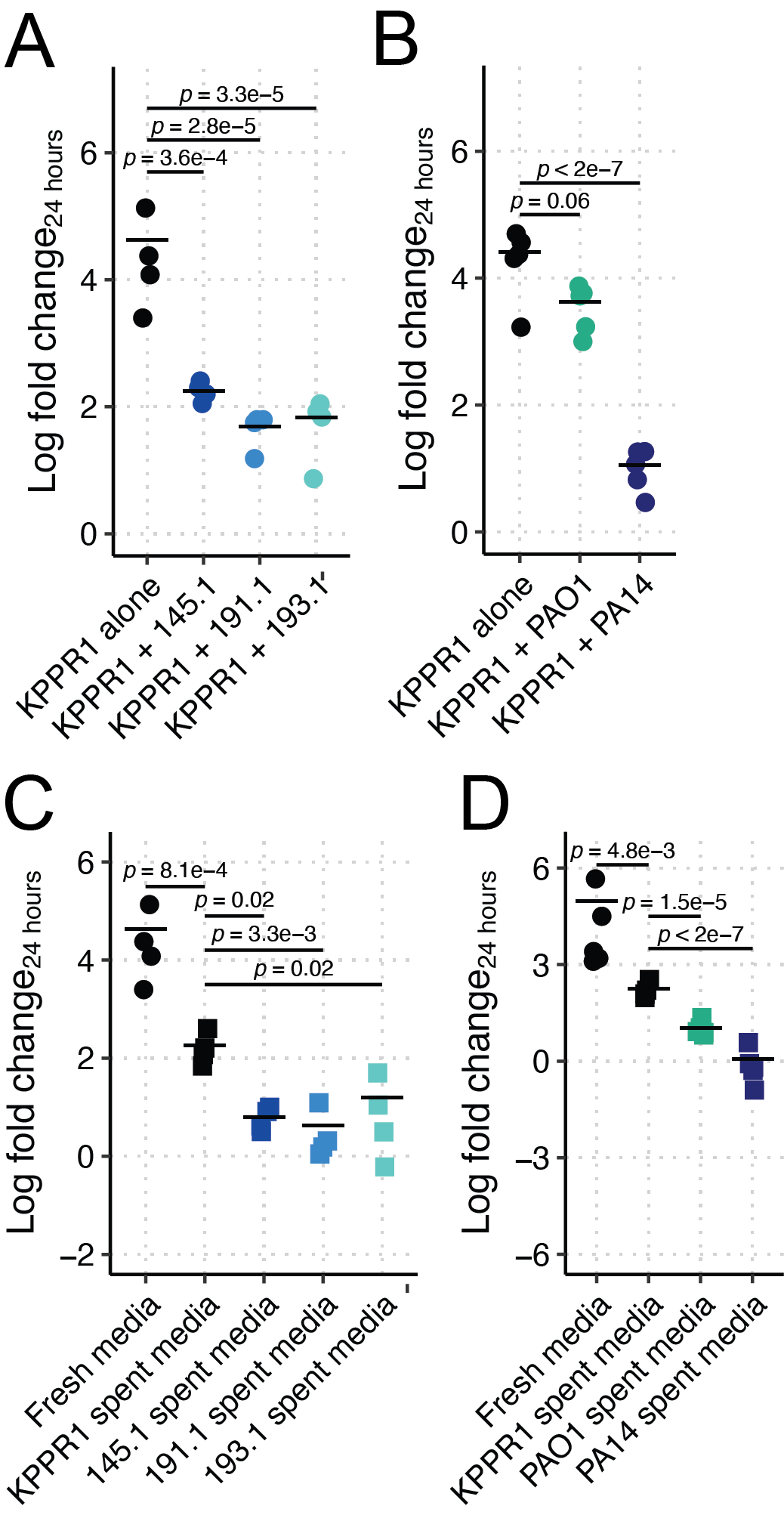

**Figure S3. Pa restricts Kp growth in M9 medium supplemented with 1.0% casamino acids in a strain-dependent, contact-independent manner.** Kp KPPR1 was grown alone or in co-culture in M9 medium supplemented with 1.0% casamino acids with mouse-derived wild Pa (**A**), PAO1, PA14 (**B**) or in filter sterilized spent media of KPPR1 or each Pa strain (**C-D**)**.** For **A-D**, “Log fold change_24 hours_” = log_10_(output KPPR1 CFU at 24 hours/input KPPR1 CFU). *p-*values represent Tukey multiple comparison correction following one-way ANOVA. Each data point is a biological replicate, and horizontal lines indicate the mean of each dataset.

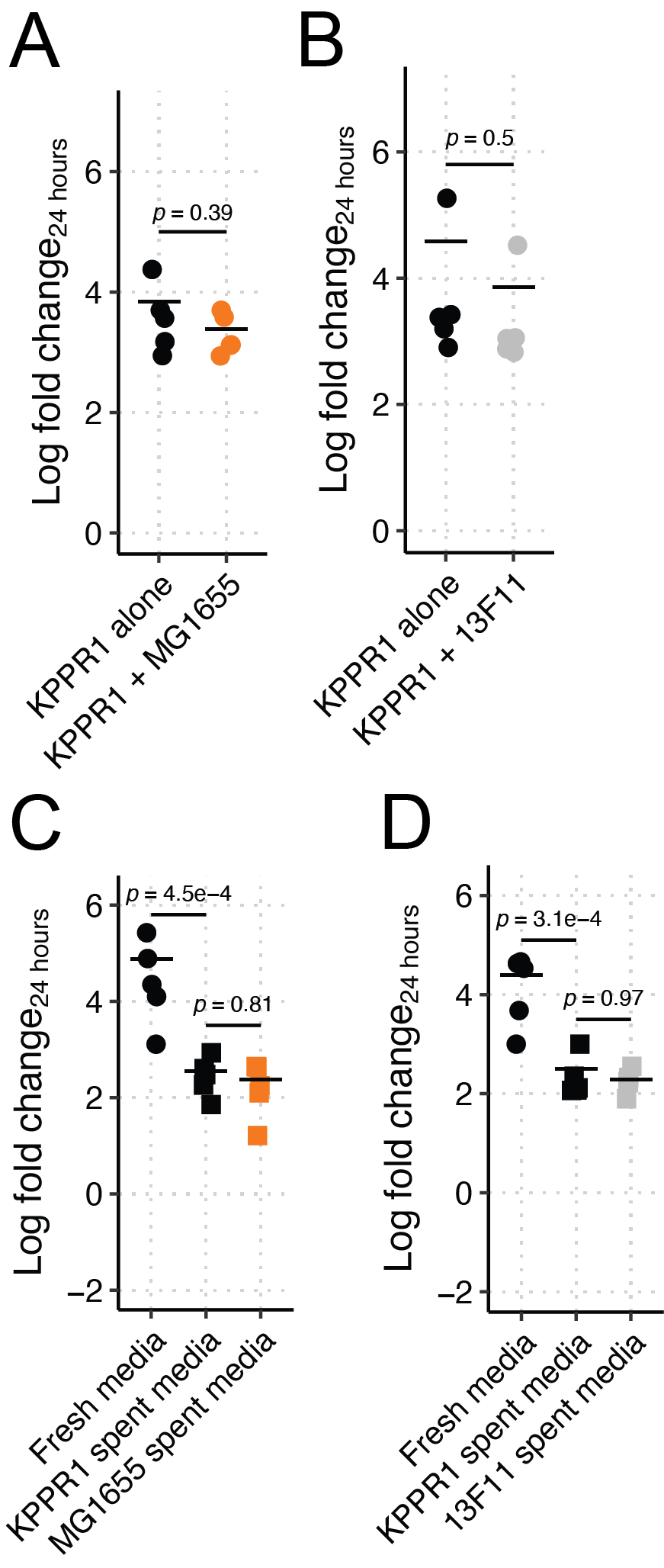

**Figure S4. Ec MG1655 and an isogenic neutral Tn mutant do not restrict KPPR1 growth in M9 medium supplemented with 1.0% casamino acids.** Kp KPPR1 was grown alone or in co-culture in M9 medium supplemented with 1.0% casamino acids with Ec MG1655 (**A**), Kp 13F11 (**B**), or in filter sterilized spent media of KPPR1, Ec MG1655 (**C**), or Kp 13F11 strain (**D**)**.** For **A-D**, “Log fold change_24 hours_” = log_10_(output KPPR1 CFU at 24 hours/input KPPR1 CFU). *p-*values represent Tukey multiple comparison correction following one-way ANOVA. Each data point is a biological replicate, and horizontal lines indicate the mean of each dataset.

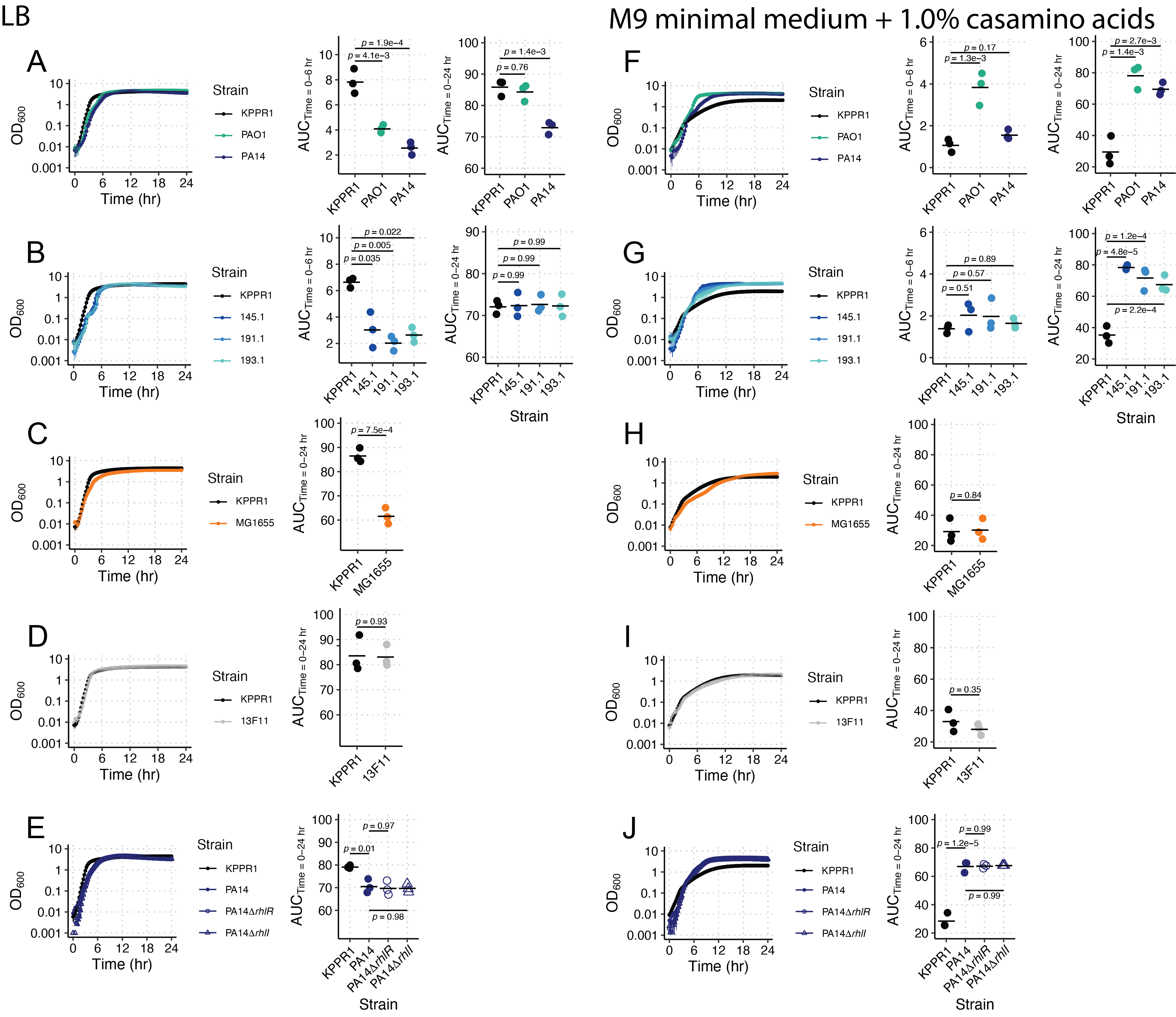

**Figure S5. Growth dynamics of select Kp, Pa, and Ec strains in LB and M9 medium supplemented with 1.0% casamino acids.** KPPR1, PAO1, PA14, Pa 145.1, Pa 191.1, Pa 193.1, MG1655, 13F11, PA14Δ*rhlR*, PA14Δ*rhlI* were grown in LB (**A-E**) or in M9 medium supplemented with 1.0% casamino acids (**F-J**). Area under the curve (AUC) analysis was used to quantify growth at early (0-6 hours, **A-B, F-G**) and late (0-24 hours, **A-J**) stages of growth. *p-*values represent Tukey multiple comparison correction following one-way ANOVA. For growth curves, each data point represents the mean, and vertical bars represent the standard error of the mean. For AUC analysis, each data point is a biological replicate, and horizontal lines indicate the mean of each dataset.

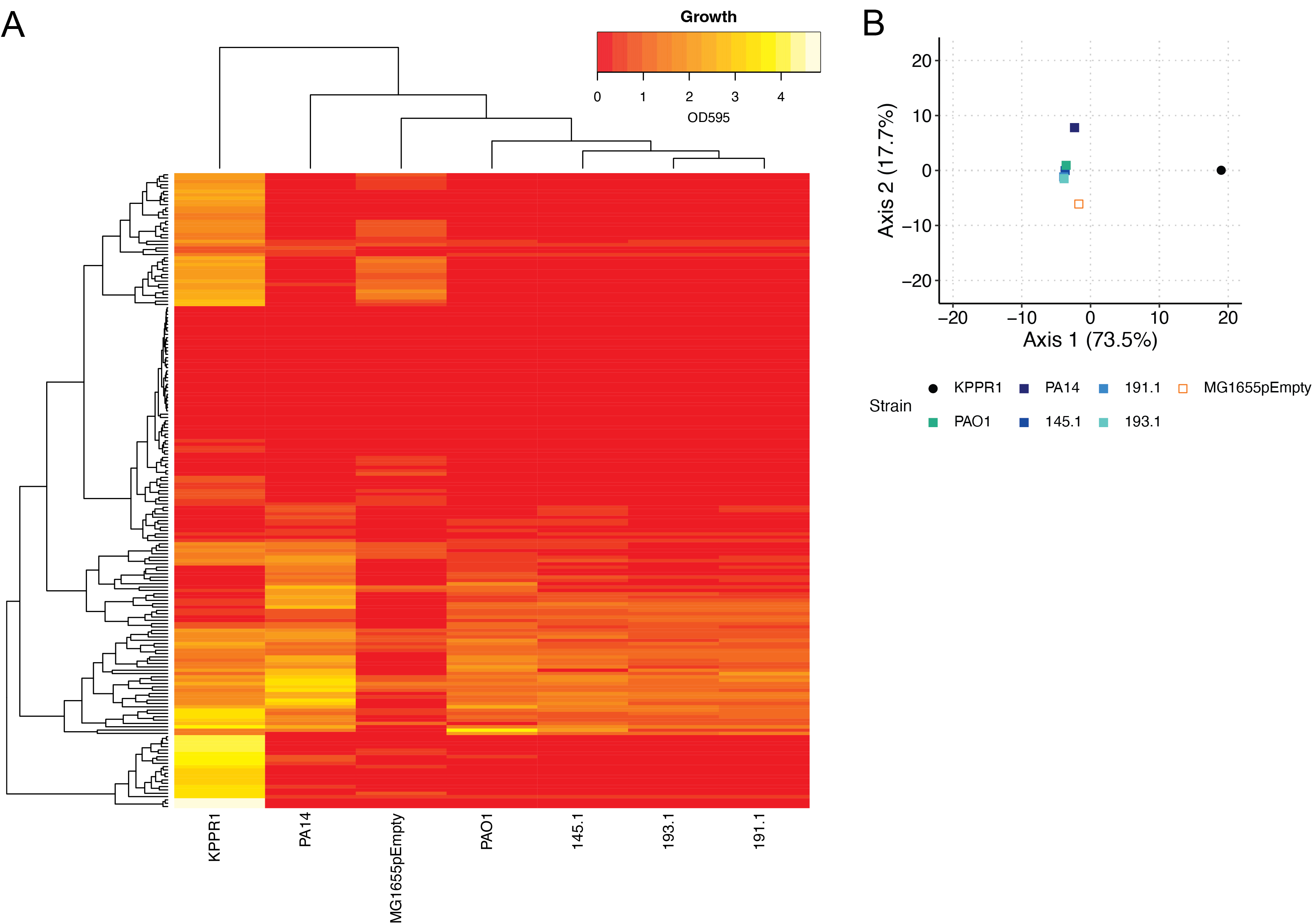

**Figure S6. WT Pa and Ec strains are less metabolically flexible than Kp.** KPPR1, PAO1, PA14, Pa 145.1, Pa 191.1, Pa 193.1, and Ec MG1655pEmpty were grown in BioLog Phenotype Microarray plates PM1 and PM2 (mean of three biological replicates displayed, each row is an individual carbon source, **A**). Euclidean distance was used to measure the dissimilarity between the growth phenotypes of each strain (**B**).

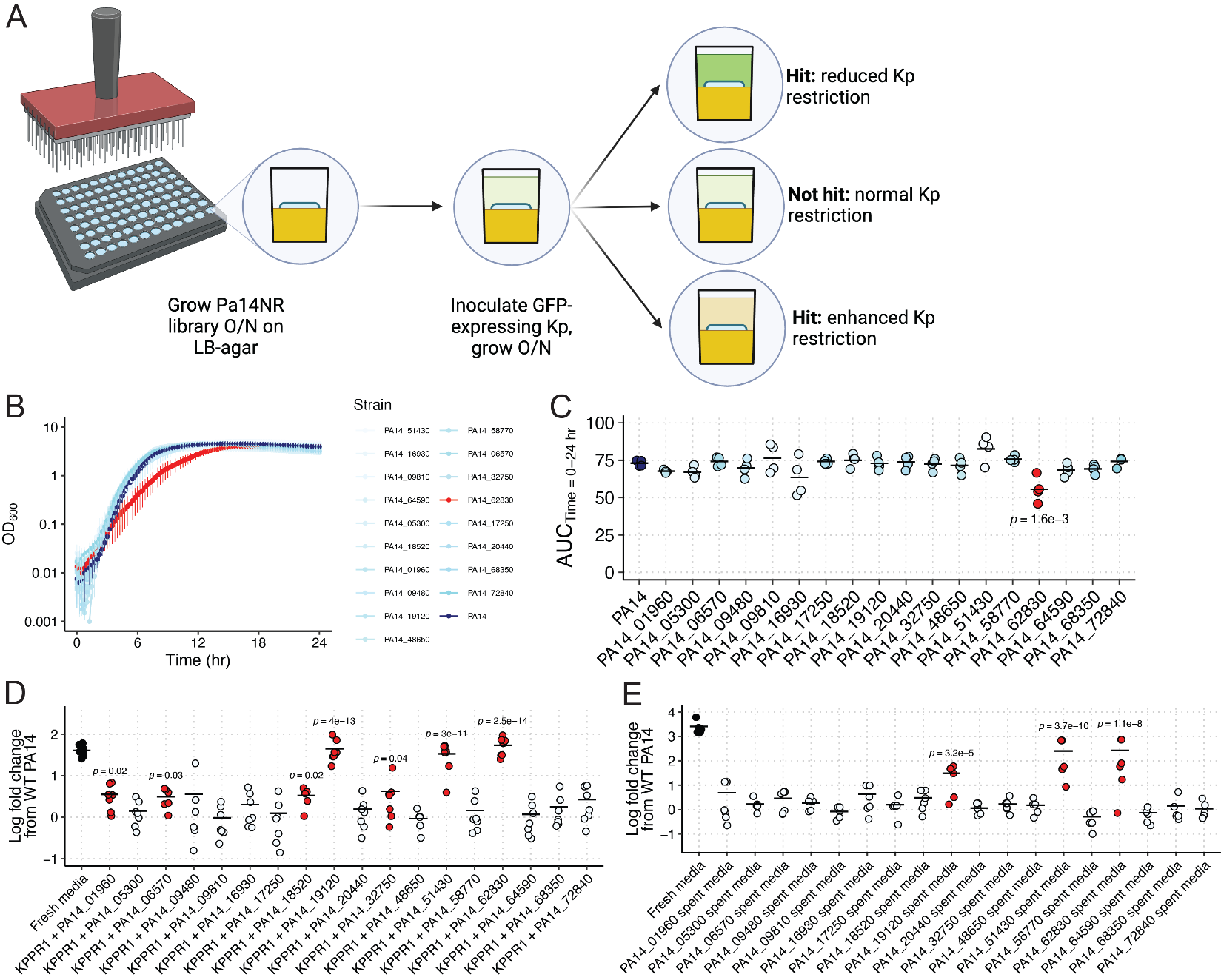

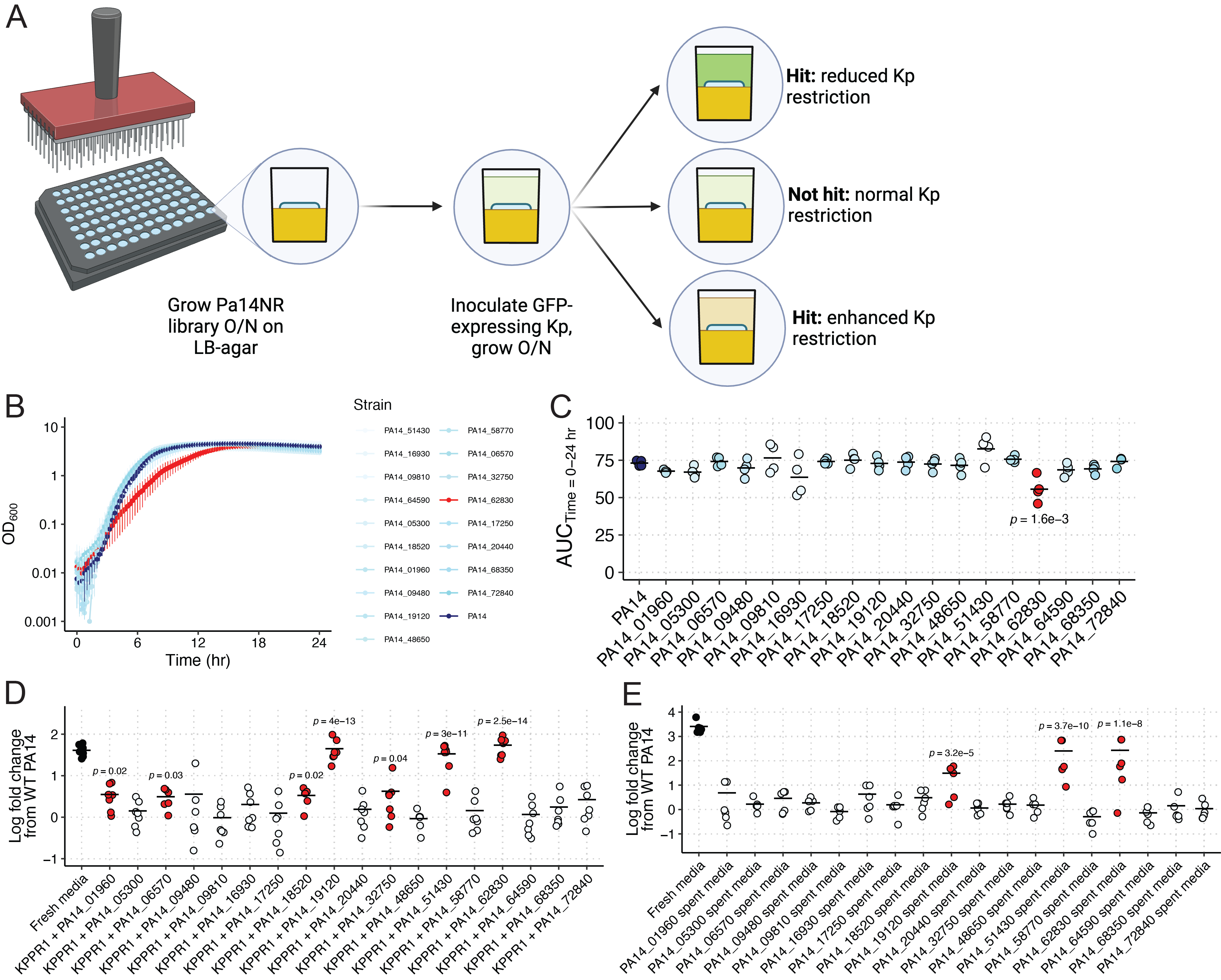

**Figure S7. PA14NR library screen and validation.** To identify candidate factors involved in Kp growth restriction, the PA14NR library was replicate plated onto LB-agar, grown overnight at 37° C, and inoculated with GFP-expressing KPPR1 (**A**). After 24 hours of co-culture, the fluorescence of each co-culture was measured, yielding 18 candidate transposon mutants, 16 of which exhibited reduced restriction, and 2 that exhibited enhanced restriction. PA14 and the 18 candidate transposon mutants identified to have a role in Kp growth restriction were grown in LB (**B**) and area under the curve (AUC) analysis was used to quantify growth (**C**). In panel **B**, each data point represents the mean, and vertical bars represent the standard error of the mean. Kp KPPR1 was grown alone or in co-culture in LB with PA14 or the 18 transposon mutants (**D**) or in filter sterilized spent media of KPPR1, PA14, or the 18 transposon mutants (**E**). For **D-E**, “Log fold change from WT PA14” = log_10_(output KPPR1 CFU at 24 hours/input KPPR1 CFU) in transposon mutant co-culture or spent media culture / log_10_(output KPPR1 CFU at 24 hours/input KPPR1 CFU) in WT PA14 co-culture or spent media culture. For **C**, *p-*values represent Tukey multiple comparison correction following one-way ANOVA, and for **D-E**, *p-*values represent one sample *t-*test from a hypothetical mean of 0. For **C-E**, each data point is a biological replicate, horizontal lines indicate the mean of each dataset, and red datasets are statistically significant from their relative comparisons.

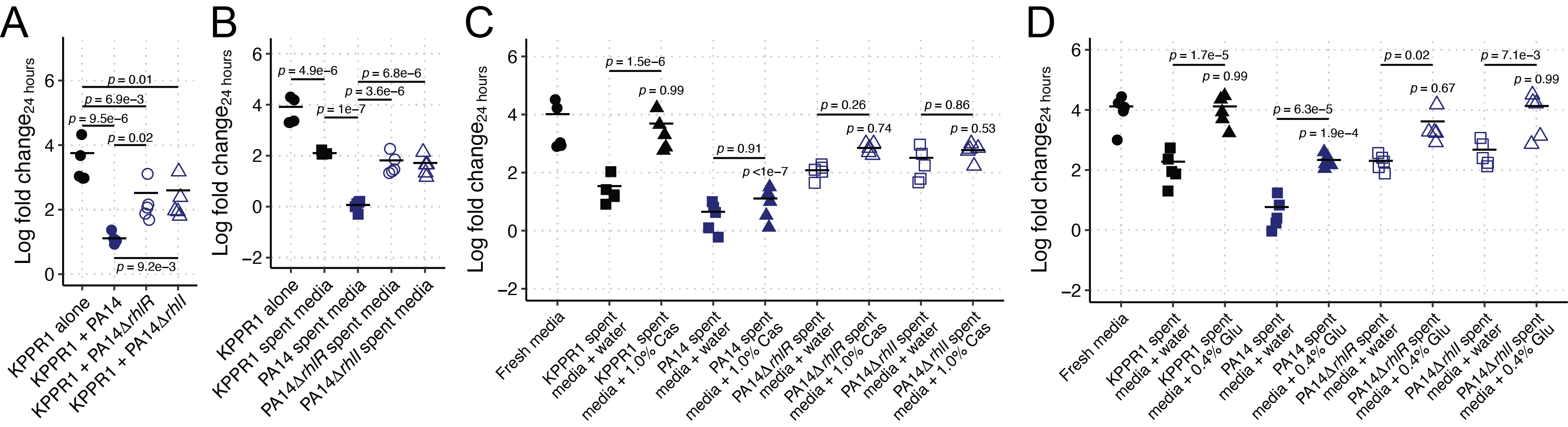

**Figure S8.** ***rhlRI* is required for Kp growth restriction by Pa in M9 medium supplemented with 1.0% casamino acids.** KPPR1 was grown alone or in co-culture in M9 medium supplemented with 1.0% casamino acids with WT PA14, PA14Δ*rhlR*, or PA14Δ*rhlI* (**A**) or in filter sterilized spent media of KPPR1 or each Pa strain (**B**), supplemented with water, 1% casamino acids (“Cas,” **C**) or 0.4% glucose (“Glu,” **D**). For **A-D**, “Log fold change_24 hours_” = log_10_(output KPPR1 CFU at 24 hours/input KPPR1 CFU). *p-*values represent Tukey multiple comparison correction following one-way ANOVA. *p-*values over columns indicate comparison to “Fresh media” condition. Each data point is a biological replicate, and horizontal lines indicate the mean of each dataset.

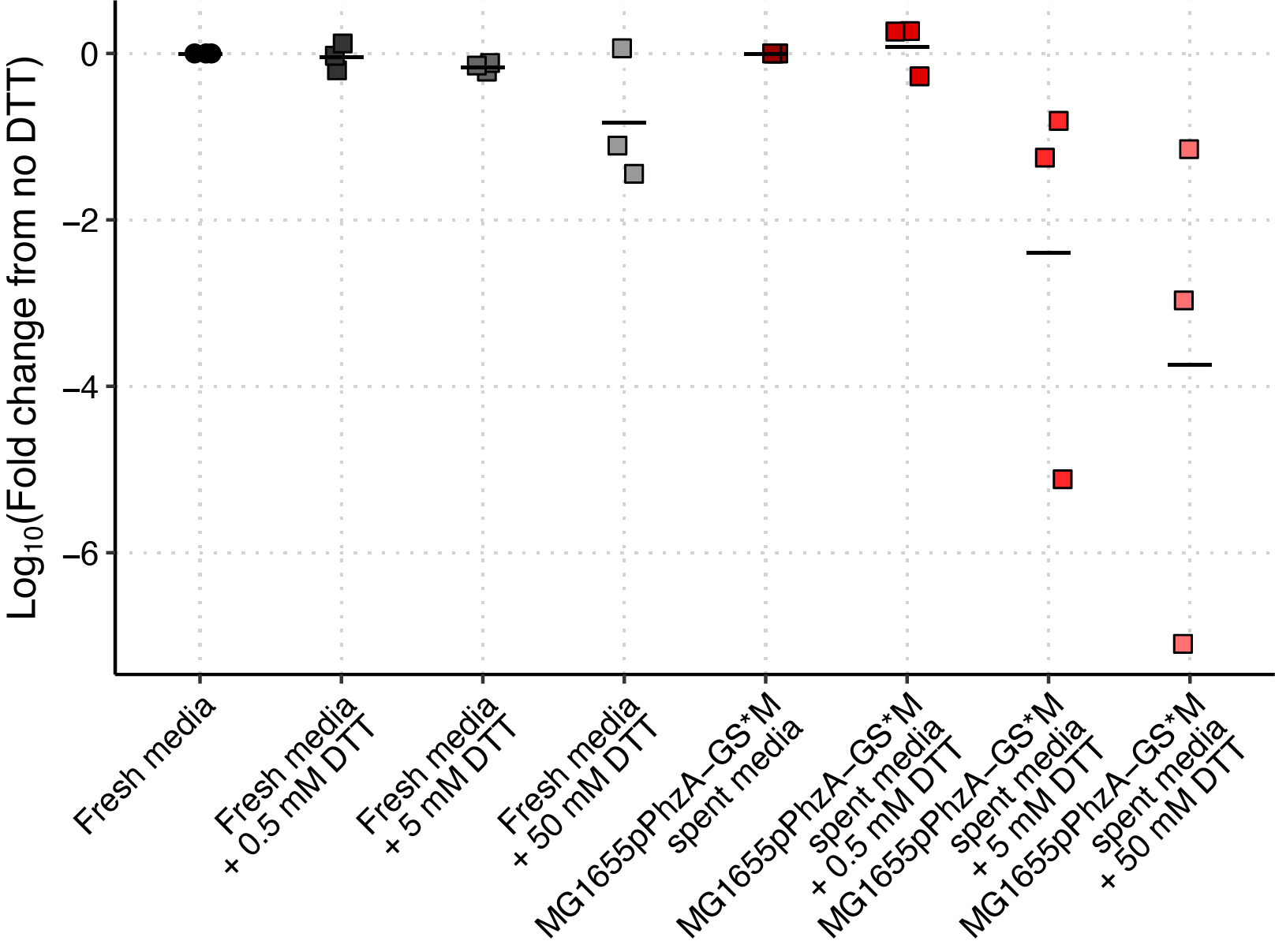

**Figure S9. 5MPCA activity is redox-dependent.** 13F11 (Kan^R^ KPPR1 variant) was grown in fresh LB broth or spent media from MG1655 constitutively expressing PYR (pPhzA-GS*M). Dithiothreitol (DTT) was titrated into both media. “Log_10_(Fold change from no DTT)” = log_10_(output 13F11 CFU at 24 hours/input 13F11 CFU) in fresh or spent media + DTT / log_10_(output 13F11 CFU at 24 hours/input 13F11 CFU) in fresh or spent media without DTT. Each data point is a biological replicate, and horizontal lines indicate the mean of each dataset.

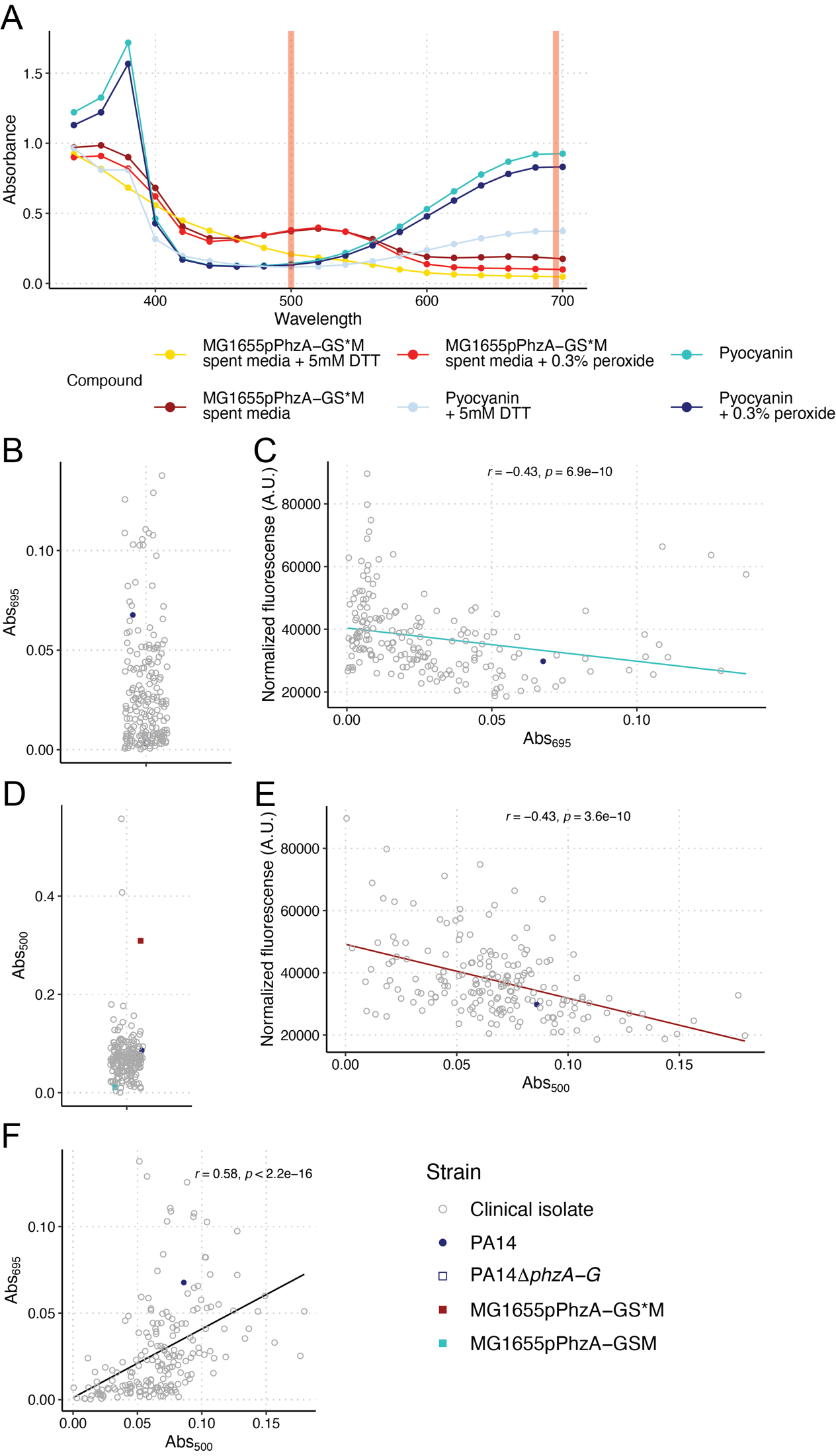

**Figure S10. Clinical Pa strain phenazine production.** The spectra of pure PYO and the spent media MG1655 constitutively expressing 5MPCA (pPhzA-GS*M) was measured under native, oxidizing (0.3% hydrogen peroxide) and reducing (5 mM dithiothreitol [DTT]) conditions (**A**) to identify wavelengths at which those phenazines can be differentiated. We determined that PYO and 5MPCA in oxidizing conditions can be differentiated at 695 and 500 nm, respectively (red vertical bars). Clinical Pa strains (N = 194), WT PA14, and PA14Δ*phzA-G* were grown in LB broth and the Abs_695_ of spent media was measured after 24 hours was measured (**B**). Abs_695_ results were correlated to growth restriction results from **Figure 5C** (Spearman correlation test, **C**). Clinical Pa strains (N = 192), WT PA14, PA14Δ*phzA-G*, and MG1655 constitutively expressing PYR (pPhzA-GS*M) and PYO (pPhzA-GSM) were grown in LB broth and the Abs_500_ of spent media was measured after 24 hours was measured following oxidation by 0.3% hydrogen peroxide (**D**). Abs_500_ results were correlated to growth restriction results from **Figure 5C** (Spearman correlation test, **E**). Abs_500_ and Abs_695_ results were correlated with one another (Spearman correlation test, **F**). Each data point represents the mean read of each culture.

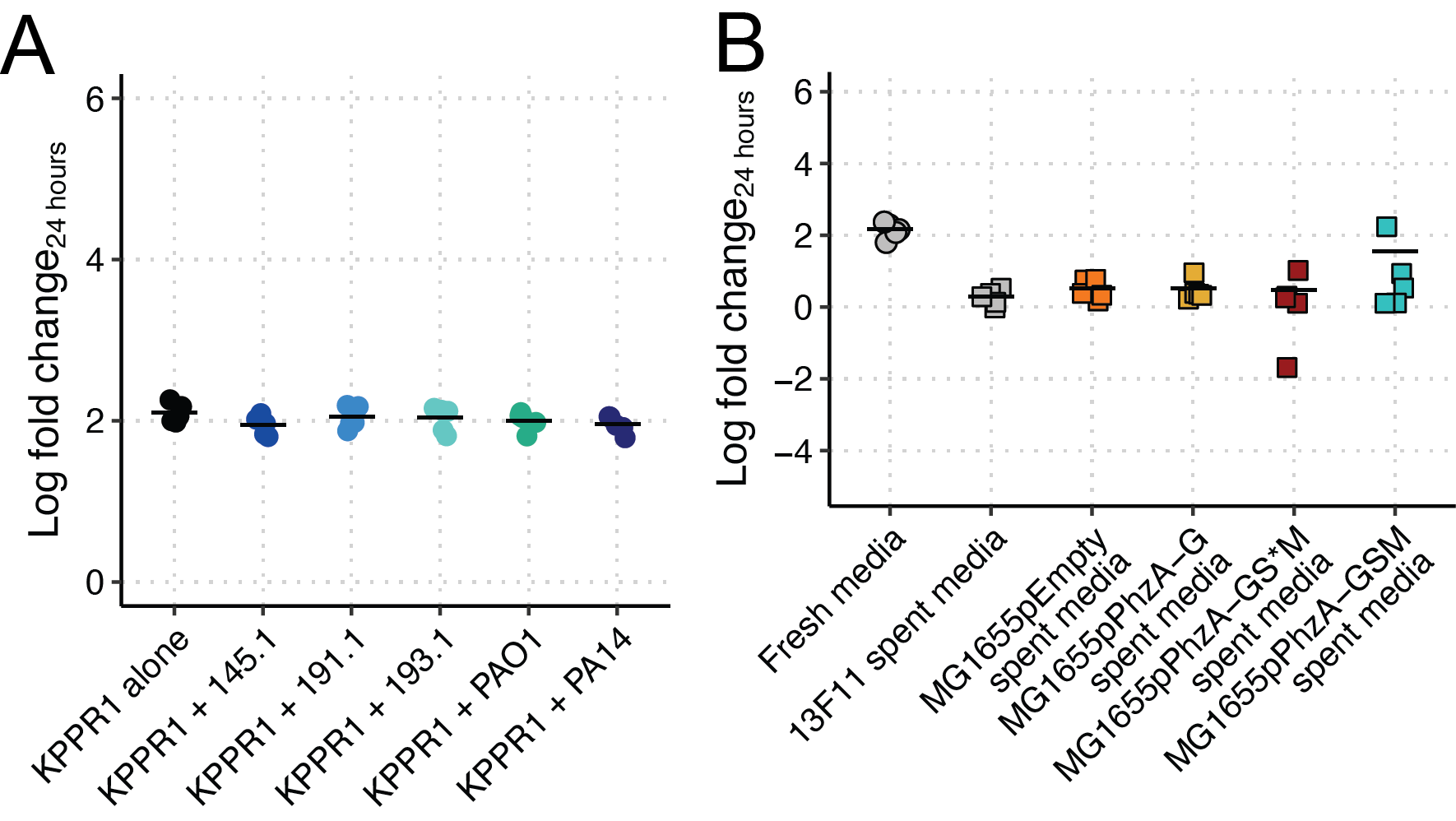

**Figure S11. Phenazines do not restrict Kp growth under anaerobic conditions.** KPPR1 was grown anaerobically alone or in co-culture in LB with mouse-derived wild Pa, PAO1, and PA14 (**A**)**.** 13F11 (Kan^R^ KPPR1 variant) was grown anaerobically in filter sterilized spent media of MG1655 containing an empty vector (pEmpty) or constitutively expressing PCA (pPhzA-G), PYR (pPhzA-GS*M), and PYO (pPhzA-GSM, **B**). For **A-B**, “Log fold change_24 hours_” = log_10_(output Kp CFU at 24 hours/input Kp CFU). Each data point is a biological replicate; horizontal lines indicate the mean of each dataset. Note that 13F11 growth in MG1655 spent media is no different than self-spent media in anerobic conditions, whereas growth is restricted in the presence of PYR and PYO in aerobic conditions (see **Figure 3F**).

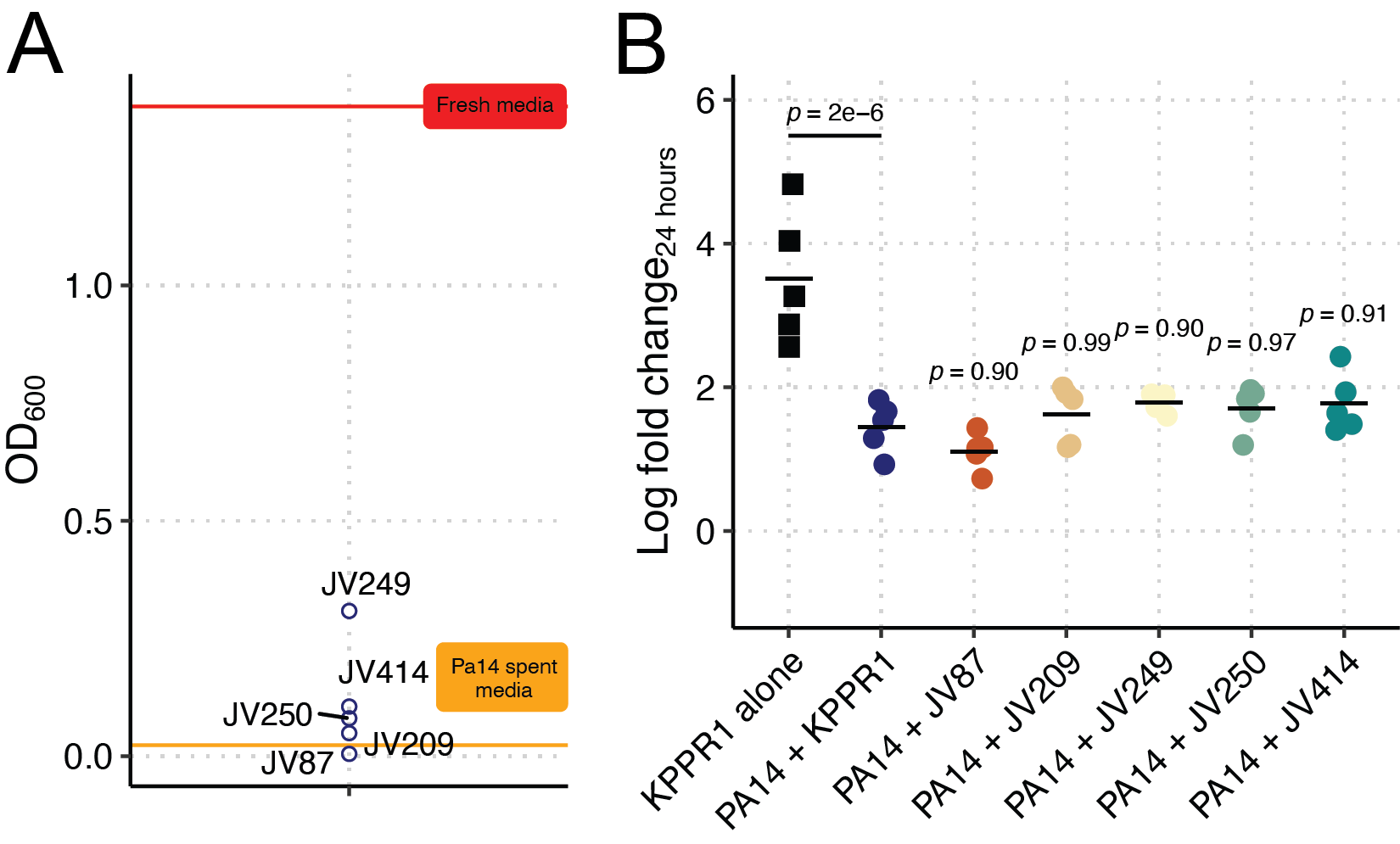

**Figure S12. Validation of clinical Kp screen results.** Clinical Kp strains (N = 6) were selected for validation (**A**). KPPR1 or select Kp strains were grown alone or co-cultured with select clinical Pa strains (**B**). For **B**, “Log fold change_24 hours_” = log_10_(output KPPR1 CFU at 24 hours/input KPPR1 CFU). *p-*values represent Tukey multiple comparison correction following one-way ANOVA compared to the “PA14 + KPPR1” condition. Each data point is a biological replicate, and horizontal lines indicate the mean of each dataset.

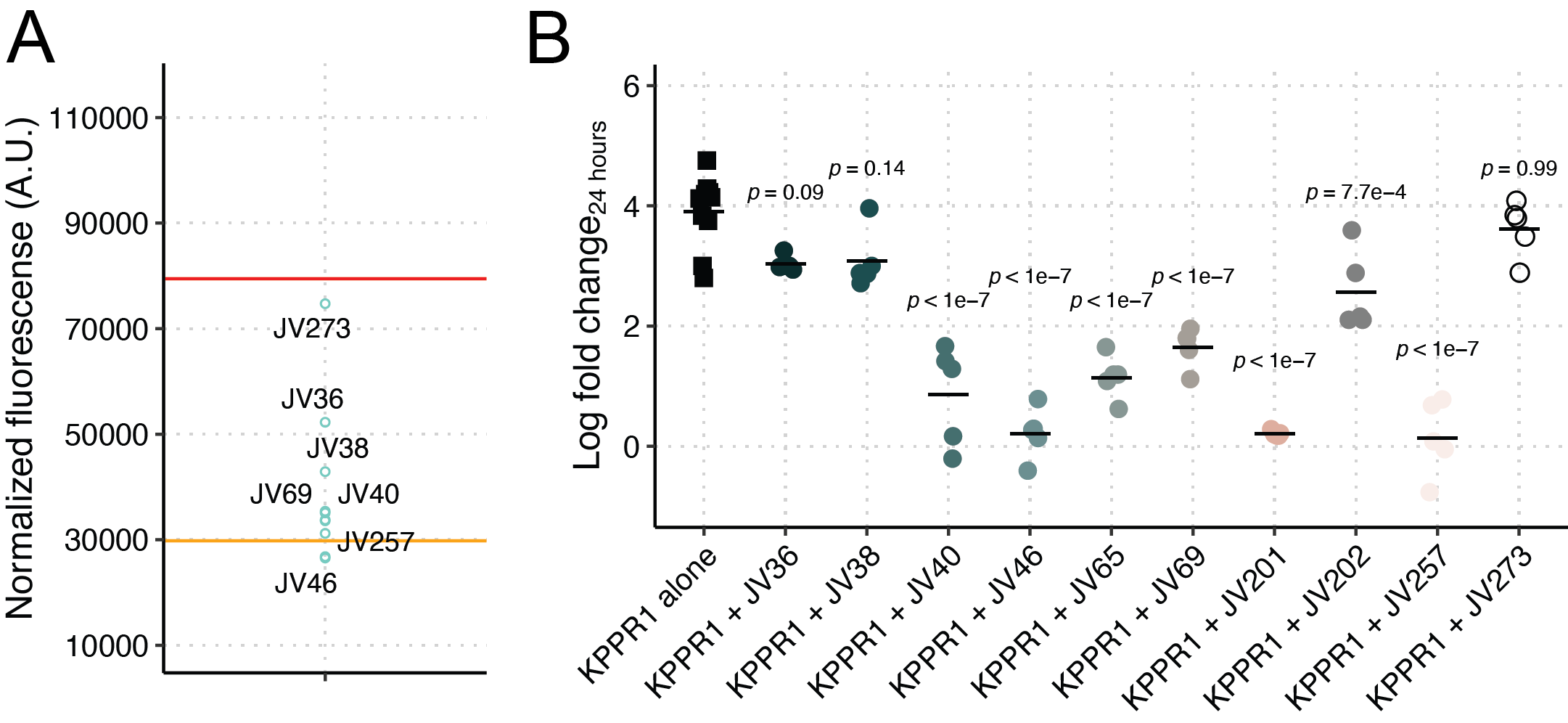

**Figure S13. Validation of clinical Pa screen results.** Clinical Pa strains (N = 10) were selected for validation (**A**). KPPR1 was grown alone or in co-culture with select clinical Pa strains (**B**). For **B**, “Log fold change_24 hours_” = log_10_(output KPPR1 CFU at 24 hours/input KPPR1 CFU). *p-*values represent Tukey multiple comparison correction following one-way ANOVA compared to the “KPPR1 alone” condition. Each data point is a biological replicate, and horizontal lines indicate the mean of each dataset.

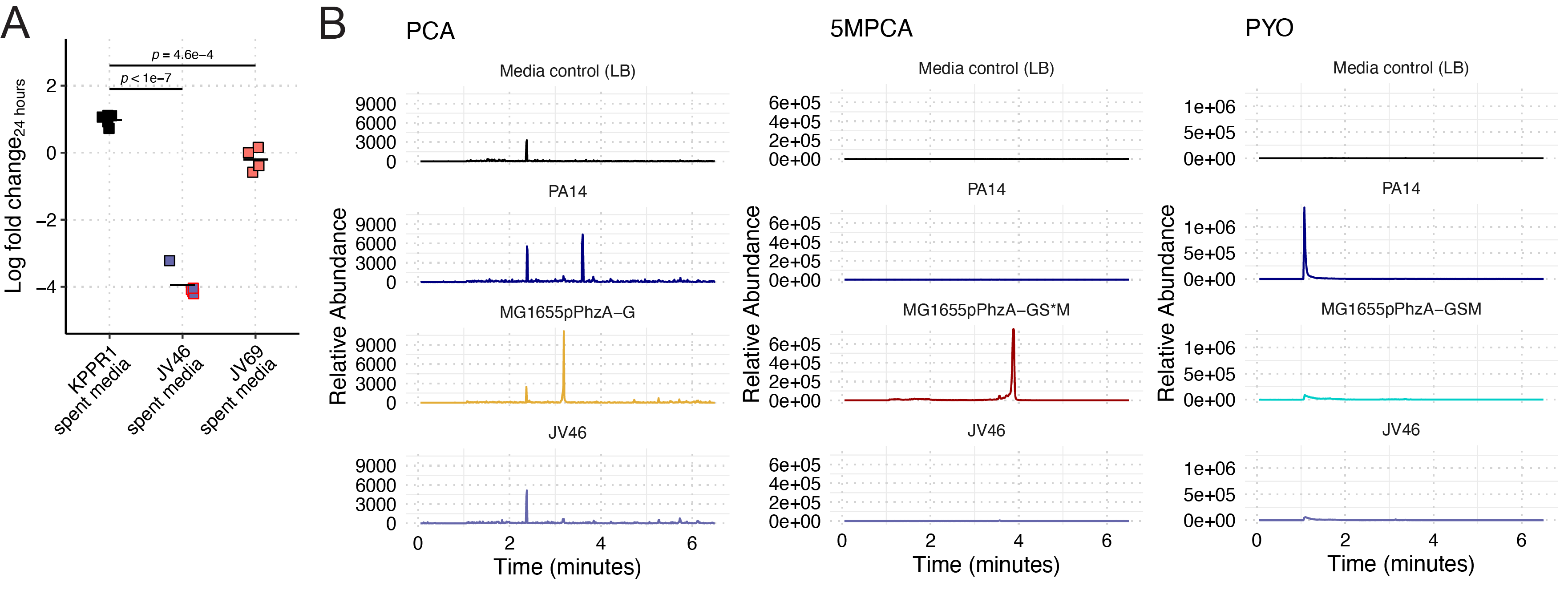

**Figure S14. Kp growth restriction in clinical Pa spent media is variable.** KPPR1 was grown in filter sterilized spent media of KPPR1, JV46, or JV69 (**A**). “Log fold change_24 hours_” = log_10_(output KPPR1 CFU at 24 hours/input KPPR1 CFU). Datapoints outlined in red are below the limit of detection (200 CFU/mL). *p-*values represent Tukey multiple comparison correction following one-way ANOVA. Each data point is a biological replicate, and horizontal lines indicate the mean of each dataset. LC-MS was used to quantify phenazine secretion from JV46 (**B**).

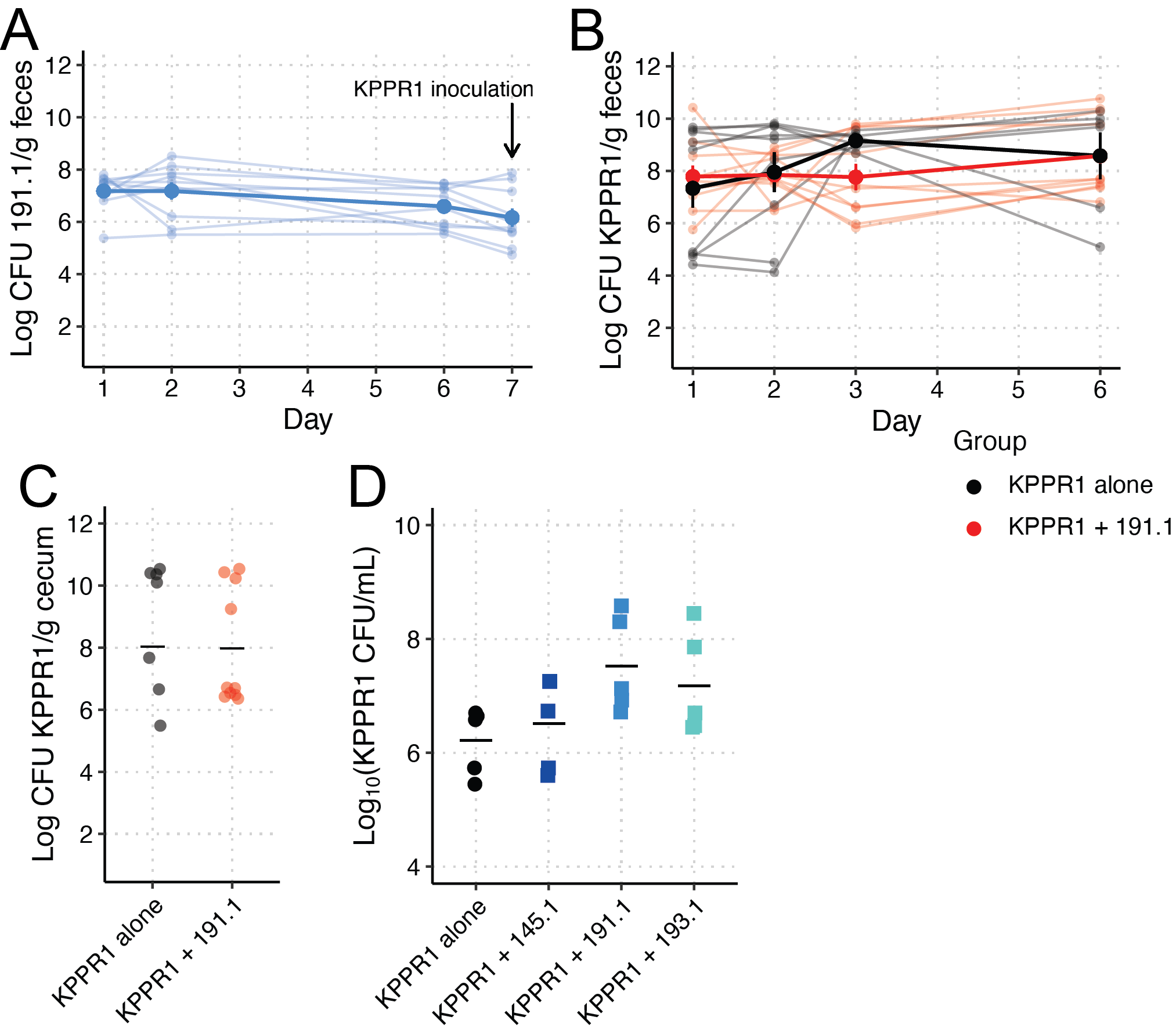

**Figure S15. Pa does not exclude Kp from the gut.** C57Bl6/J mice were treated for 4 days with 0.5 g/L ampicillin, then orally gavaged with ~10^7^ CFU 191.1. Seven days post-191.1 colonization, mice were orally gavaged a second time with ~10^8^ CFU KPPR1 (N = 10). Antibiotic-treated KPPR1 mono-colonized mice served as a control (N = 10). 191.1 fecal loads were monitored post-colonization (**A**), and KPPR1 fecal (**B**) and cecal loads (**C**) were measured were monitored post-colonization (after 7 days 191.1 colonization) or at the end of the experiment, respectively. Large intestinal contents were from C57Bl6/J (no antibiotic treatment, N = 5) and resuspended in sterile PBS. ~5x10^7^ CFU KPPR1 alone or an equal ratio of KPPR1 with each wild Pa, was inoculated into large intestinal contents and grown anaerobically. KPPR1 density was measured at 48 hours (**D**).

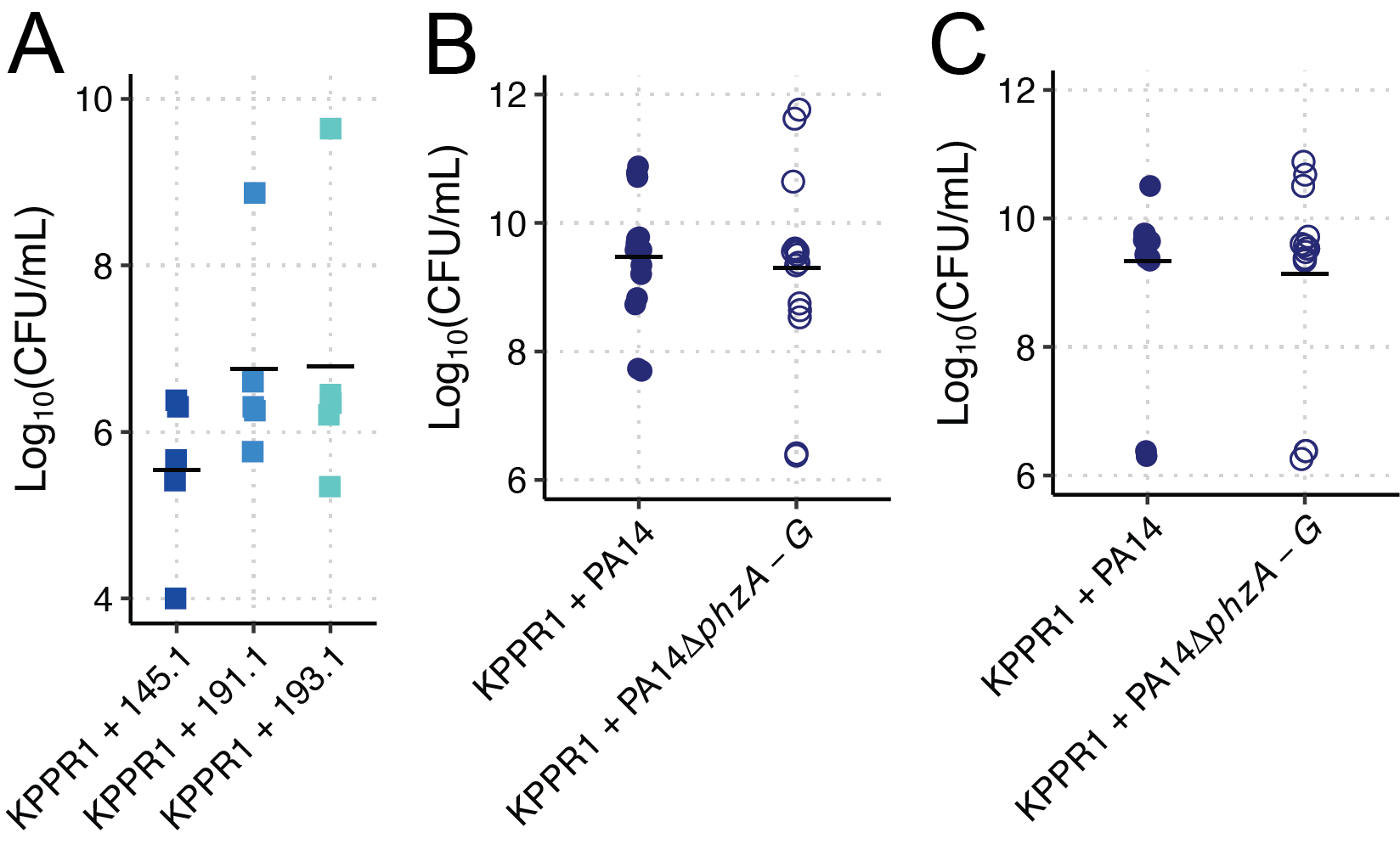

**Figure S16. Pa is viable in *ex vivo* tissues.** Pa density was measured in large intestinal contents (**A**), BALF (**B**), and bladder homogenate (**C**) from experiments presented in **Figures S15A**, **6C**, and **6D**, respectively.

**Table S1. Clinical Pa isolation sources**

| **Strain** | **Isolation source (verbatim from clinical lab)** | **Clinical code** |
| --- | --- | --- |
| JV15 | heel wound | wound |
| JV16 | abdominal wound | wound |
| JV17 | urine | urine |
| JV18 | left eye | other |
| JV19 | left native bronchus | respiratory |
| JV20 | knee drainage | other |
| JV21 | left leg wound | wound |
| JV22 | right ear drainage | other |
| JV23 | wound (back) | wound |
| JV24 | urine | urine |
| JV26 | left native bronchus isolate 2 | respiratory |
| JV27 | heel wound | wound |
| JV28 | left foot wound | wound |
| JV29 | leg wound | wound |
| JV30 | references ast | other |
| JV31 | urine | urine |
| JV32 | bal fluid | respiratory |
| JV33 | blood | blood |
| JV34 | sternum wound | wound |
| JV35 | heart tissue | other |
| JV36 | nasal wound | wound |
| JV38 | urine | urine |
| JV39 | bal fluid | respiratory |
| JV40 | cranial tissue | other |
| JV41 | urine | urine |
| JV42 | right leg wound | wound |
| JV43 | left ankle wound | wound |
| JV44 | blood | blood |
| JV45 | ankle wound | wound |
| JV46 | blood | blood |
| JV47 | urine | urine |
| JV48 | sputum | respiratory |
| JV49 | sputum | respiratory |
| JV50 | cf throat culture | respiratory |
| JV51 | tracheal aspirate | respiratory |
| JV52 | tracheal aspirate | respiratory |
| JV53 | sputum | respiratory |
| JV54 | right foot | other |
| JV55 | sputum | respiratory |
| JV56 | bal fluid | respiratory |
| JV57 | tracheal aspirate | respiratory |
| JV58 | burn isolate 2 | other |
| JV59 | burn isolate 1 | other |
| JV60 | blood | blood |
| JV61 | urine | urine |
| JV62 | urine | urine |
| JV63 | blood | blood |
| JV64 | right leg | other |
| JV65 | urine | urine |
| JV66 | urine | urine |
| JV67 | urine | urine |
| JV68 | urine | urine |
| JV69 | urine | urine |
| JV70 | urine | urine |
| JV71 | urine | urine |
| JV73 | urine | urine |
| JV193 | urine | urine |
| JV194 | urine | urine |
| JV195 | urine | urine |
| JV196 | urine | urine |
| JV197 | groin abscess | other |
| JV198 | wound | wound |
| JV199 | left ear | other |
| JV200 | gluteal wound | wound |
| JV201 | ear | other |
| JV202 | wound | wound |
| JV203 | left ear | other |
| JV204 | left leg wound | wound |
| JV257 | right leg | other |
| JV258 | ear drainage | other |
| JV259 | wound | wound |
| JV260 | left leg | other |
| JV261 | urine | urine |
| JV262 | discharge | other |
| JV263 | leg wound | wound |
| JV264 | right ear | other |
| JV265 | left leg wound | wound |
| JV266 | urine | urine |
| JV267 | urine | urine |
| JV268 | urine | urine |
| JV269 | urine | urine |
| JV270 | vaginal discharge | other |
| JV271 | decubitus ulcer | other |
| JV272 | hip wound | wound |
| JV273 | pelvic abscess | other |
| JV274 | bal fluid | respiratory |
| JV275 | urine | urine |
| JV276 | right leg | other |
| JV287 | urine | urine |
| JV288 | buttock tissue | other |
| JV289 | trach aspirate | respiratory |
| JV290 | knee tissue | other |
| JV291 | bal fluid | respiratory |
| JV292 | cf patient | other |
| JV294 | calf tissue | other |
| JV296 | tissue | other |
| JV297 | abscess | other |
| JV298 | sputum | respiratory |
| JV299 | hardware | other |
| JV301 | urine | urine |
| JV302 | leg tissue | other |
| JV303 | trach aspirate | respiratory |
| JV304 | bal fluid | respiratory |
| JV305 | bal fluid | respiratory |
| JV306 | sputum | respiratory |
| JV307 | bal fluid | respiratory |
| JV308 | cf patient throat | respiratory |
| JV309 | cf patient throat | respiratory |
| JV310 | bal fluid | respiratory |
| JV311 | trach aspirate | respiratory |
| JV312 | sputum | respiratory |
| JV313 | cf patient throat | respiratory |
| JV314 | trach aspirate | respiratory |
| JV315 | bal fluid rll | respiratory |
| JV316 | urine | urine |
| JV317 | urine | urine |
| JV318 | urine | urine |
| JV319 | bone | other |
| JV320 | urine | urine |
| JV321 | cf patient throat | respiratory |
| JV322 | urine | urine |
| JV323 | urine | urine |
| JV324 | wound | wound |
| JV325 | sputum | respiratory |
| JV326 | right foot wound | wound |
| JV327 | bal fluid | respiratory |
| JV328 | respiratory | respiratory |
| JV329 | trach aspirate | respiratory |
| JV330 | sputum | respiratory |
| JV331 | sputum | respiratory |
| JV332 | bal fluid | respiratory |
| JV333 | urine | urine |
| JV334 | urine | urine |
| JV335 | urine | urine |
| JV336 | urine | urine |
| JV337 | stool | other |
| JV340 | urine | urine |
| JV341 | urine | urine |
| JV342 | right foot bone | other |
| JV343 | urine | urine |
| JV344 | urine | urine |
| JV345 | urine | urine |
| JV346 | urine | urine |
| JV347 | urine | urine |
| JV348 | ear drainage | other |
| JV349 | urine | urine |
| JV350 | drainage | other |
| JV351 | chest wound | wound |
| JV352 | cheek drainage | other |
| JV353 | cath site drainage | other |
| JV354 | sacrum drainage | other |
| JV355 | foot | other |
| JV356 | right leg wound | wound |
| JV357 | urine | urine |
| JV358 | abdomen | other |
| JV359 | urine | urine |
| JV360 | abscess left leg | other |
| JV361 | urine | urine |
| JV362 | urine | urine |
| JV363 | right foot wound | wound |
| JV364 | left foot wound | wound |
| JV365 | left leg wound | wound |
| JV366 | bal fluid | respiratory |
| JV367 | tracheal aspirate | respiratory |
| JV368 | sputum | respiratory |
| JV369 | sputum | respiratory |
| JV370 | tracheal aspirate | respiratory |
| JV371 | right leg wound | wound |
| JV372 | bal fluid | respiratory |
| JV373 | bal fluid | respiratory |
| JV374 | cf patient | other |
| JV375 | left leg drainage | other |
| JV376 | bronch wash | respiratory |
| JV377 | right leg wound | wound |
| JV378 | left foot | other |
| JV379 | urine | urine |
| JV380 | ear drainage | other |
| JV381 | bal fluid | respiratory |
| JV382 | bal fluid | respiratory |
| JV384 | sputum | respiratory |
| JV391 | sputum | respiratory |
| JV392 | bal fluid | respiratory |
| JV393 | urine | urine |
| JV394 | back drainage | other |
| JV395 | urine | urine |
| JV415 | abdominal fluid | other |
| JV416 | blood-pediatric | blood |
| JV417 | urine-cath | urine |
| JV418 | respiratory | respiratory |
| JV419 | urine | urine |
| JV420 | urine-cath | urine |
| JV421 | right leg | other |
| JV422 | urostomy | urine |
| JV423 | right leg tissue | other |
| JV424 | urine | urine |
